# ACE-engineered human iPSC-derived neutrophils enhance host defense against multidrug-resistant bacterial infections

**DOI:** 10.64898/2026.11.09.765250

**Authors:** Tomohiro Shibata, Shabir A. Bhat, Suguru Saito, Tahir B. Dar, Binod GC, Ellen A. Bernstein, Noor Khan, Dhruv Sareen, Helen S. Goodridge, Warren G. Tourtellotte, Kenneth E. Bernstein, Zakir Khan

**Affiliations:** Department of Biomedical Sciences, Cedars-Sinai Medical Center, Los Angeles, CA, United States; Department of Pathology and Laboratory Medicine, Cedars-Sinai Medical Center, Los Angeles, CA, United States; Department of Pharmacology, Yokohama City University School of Medicine, Yokohama, Japan; Department of Pathology and Laboratory Medicine, David Geffen School of Medicine at University of California Los Angeles (UCLA), CA, United States; Laboratory of Integrative Metabolic Regulation, Institute for Molecular and Cellular Regulation, Gunma University, Maebashi, Gunma 3718512, Japan; Department of Medicine, Division of Hematology and Oncology, David Geffen School of Medicine at University of California, Los Angeles (UCLA), Los Angeles, CA, United States; The Board of Governors Regenerative Medicine Institute, Cedars-Sinai Medical Center, Los Angeles, CA, United States; Cedars-Sinai Cancer, Cedars-Sinai Medical Center, Los Angeles, CA, United States; Department of Neurology and Neurosurgery, Cedars-Sinai Medical Center, Los Angeles, CA, United States; Institute for Myeloma & Bone Cancer Research (IMBCR), West Hollywood, CA, United States

**Keywords:** iPSCs, Neutrophils, Angiotensin-Converting Enzyme (ACE), Granulocyte Transfusion, Multidrug-Resistant Infections, Neutropenia

## Abstract

Severe neutropenia predisposes patients to life-threatening infections, while donor-dependent granulocyte transfusions are limited by availability and variable cell quality. Here, we engineered human induced pluripotent stem cells (iPSCs) to constitutively or inducibly express angiotensin-converting enzyme (ACE) and differentiated them into functional ACE-engineered neutrophils (ACE-iNeu). ACE expression enhanced granulocytic maturation, reactive oxygen species generation, inflammatory cytokine production, and antimicrobial signaling. ACE-iNeu exhibited enhanced bactericidal activity against methicillin-resistant *Staphylococcus aureus* and multidrug-resistant *Pseudomonas aeruginosa*. Following adoptive transfer into neutropenic mice, ACE-iNeu significantly reduced circulating and tissue bacterial burdens. Integrated metabolomic and transcriptomic analyses revealed ACE-associated alterations in adenine nucleotide abundance and developmental and inflammatory pathways. Together, these findings establish ACE engineering as a strategy to enhance the production and antimicrobial function of human iPSC-derived neutrophils and support further preclinical development of engineered granulocyte therapies for severe drug-resistant infections.

**Highlights:**

- ACE enhances human iPSC-derived neutrophil maturation and antimicrobial effector functions.
- ACE-engineered iPSC-derived neutrophils improve clearance of multidrug-resistant bacterial infections in vivo.

## Introduction

Despite advances in modern antimicrobials and supportive care, bacterial and fungal infections remain major complications in patients with prolonged disease- or therapy-induced neutropenia.^1^ For instance, myelosuppressive chemotherapy are at risk of developing chemotherapy-induced neutropenia (CIN).^2,3^ Neutropenia predisposes patients to potentially life-threatening complications, including febrile neutropenia, hospitalization, and increased mortality.^4^ The clinical introduction of granulocyte colony-stimulating factors (G-CSFs) has significantly reduced these complications by stimulating neutrophil production and maturation in patients undergoing chemotherapy.^2,5^ Several emerging therapies with mechanisms of action different from those of standard G-CSF agents have recently been approved and have the potential to improve the management of CIN.^2^ However, G-CSF induces bone pain, and continued vulnerability to infection in the first week after chemotherapy remains problematic.^2,6^ In addition, antibiotic resistance is a major global health threat with an estimated 1.27 million deaths annually, and it continues to increase in scope and severity every year.^7^ There has been increasing demand for donated granulocyte transfusions to treat or prevent severe drug-resistant infections in patients who lack their own functional granulocytes.^8^ Therefore, new approaches are required to control neutropenia and severe infections.

Angiotensin-converting enzyme (ACE) plays a critical role in blood pressure regulation through the conversion of angiotensin I to angiotensin II (Ang II).^9,10^ However, ACE is widely expressed across multiple tissues and functions as a pleiotropic peptidase capable of cleaving numerous substrates, which can influence many physiological processes.^9,10^ Our laboratory has been investigating the novel roles of ACE in myelomonocytic cells, particularly its contribution to neutrophil- and macrophage-mediated immune responses. These studies have demonstrated that ACE is a key regulator of cellular metabolism and, consequently, modulates host responses to a wide range of immune challenges, including tumor progression and bacterial infection.^11–14^ In murine models, ACE deficiency significantly impaired neutrophil antibacterial activity, whereas increased ACE expression enhanced neutrophil effector functions both in vitro and in vivo. Mechanistically, increased ACE expression was associated with higher leukotriene B4 production, longer neutrophil survival, and greater antibacterial activity.^11,12^ Consistent with these findings, we subsequently demonstrated that the ACE-dependent mechanisms regulating antimicrobial function are conserved in primary human neutrophils. ACE catalytic activity blocking using ramipril, in healthy volunteers significantly reduced neutrophil reactive oxygen species (ROS) production and impaired bactericidal activity.^12^ Collectively, these studies established ACE as a critical regulator of neutrophil innate immune function and suggested that enhancing ACE expression may represent a novel therapeutic strategy for strengthening host defense against severe bacterial infections.

To further investigate the role of ACE in human neutrophils and explore its potential application in granulocyte transfusion therapy, we generated ACE-expressing human neutrophils from genetically engineered induced pluripotent stem cells (iPSCs) and evaluated their therapeutic efficacy in models of neutropenia and antibiotic-resistant bacterial infection.

## Results

### Generation of constitutive and inducible ACE-expressing iPSC lines

To investigate the role of ACE in human granulocytes, we generated iPSC lines with targeted transgene integration at the AAVS1 safe-harbor locus. For constitutive expression, we constructed an AAVS1-CMV-hACE donor plasmid containing human ACE cDNA driven by the CMV promoter and flanked by left and right homology arms for CRISPR-Cas9-mediated knock-in (**Figure 1A**). Co-transfection of a wild-type fibroblast-derived iPSC line (CS88iCTR-nxx) with the donor construct and a Cas9 expression vector generated an ACE-overexpressing iPSC line (ACE-iPSC). In parallel, a control line (Cnt-iPSC) was established using an identical donor construct lacking ACE cDNA. Immunoblot analysis confirmed robust constitutive ACE expression in ACE-iPSC cells, with approximately 40-fold higher ACE protein levels than in control cells (**Figure 1B**).

**Figure 1.**
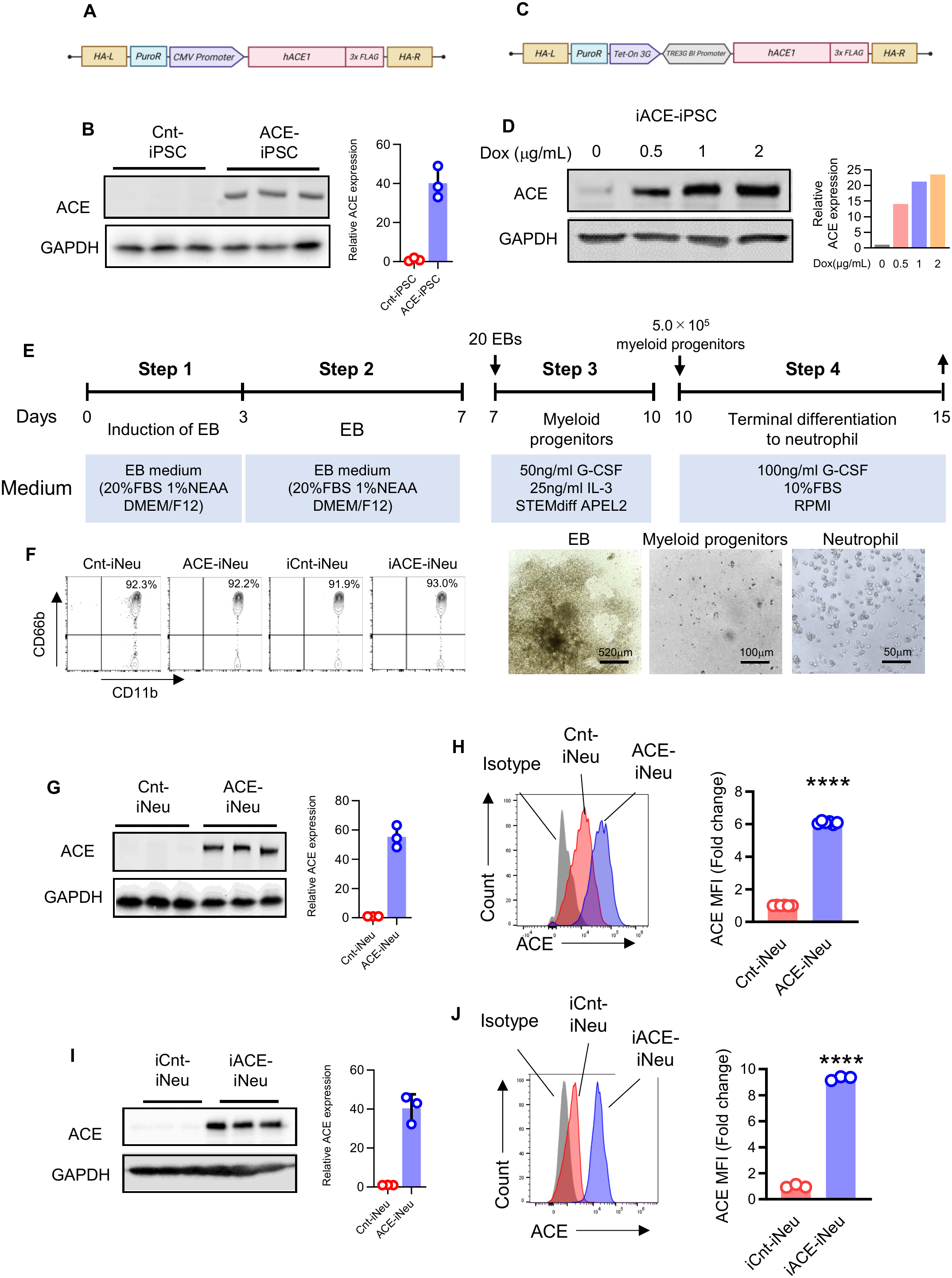
Generation of constitutive and inducible ACE-expressing human iPSC lines and differentiation into ACE-engineered neutrophils. (A) Schematic of the AAVS1-CMV-hACE construct used to generate an iPSC line with constitutive ACE expression driven by the CMV promoter (ACE-iPSC). (B) Immunoblot analysis of ACE expression in ACE-iPSCs and control iPSCs (Cnt-iPSCs) carrying an empty AAVS1-targeting vector lacking hACE cDNA. (C) Schematic of the AAVS1-Tet-On-hACE construct used to generate an iPSC line with doxycycline (Dox)-inducible ACE expression (iACE-iPSCs). (D) Immunoblot analysis of ACE expression in iACE-iPSCs cells following Dox induction. (E) Schematic of the directed differentiation protocol from iPSCs to mature neutrophils (iNeu), with representative bright-field images of embryoid bodies (EBs, day 7), myeloid progenitors (day 10), and mature neutrophils (day 15). (F) Representative flow cytometry analysis of day 15 iNeu generated from ACE-iPSC, Cnt-iPSC, iACE-iPSC and iCnt-iPSC lines. (G, H) Assessment of ACE expression in Cnt-iNeu and ACE-iNeu by immunoblotting (G) and flow cytometry (H). (I, J) Assessment of ACE expression in iCnt-iNeu and iACE-iNeu by immunoblotting (I) and flow cytometry (J). Flow cytometry data are presented as representative histograms (left) and quantification of mean fluorescence intensity (MFI; right). Data are presented as mean ± SEM. Statistical significance was determined using a two-tailed unpaired Student’s *t*-test. ****p < 0.0001.

To enable temporal regulation of ACE expression, we next generated a doxycycline (Dox)-inducible iPSC line using a Tet-On-hACE construct targeted to the same AAVS1 locus (**Figure 1C**). The donor cassette contained ACE cDNA under the control of a tetracycline-responsive promoter together with the Tet-On regulatory system, allowing inducible transgene expression following Dox administration. Correctly targeted clones (iACE-iPSC) displayed Dox-dependent ACE induction, with ACE expression increasing in a dose-responsive manner and reaching maximal levels at 1 μg/ml Dox (**Figure 1D**). These engineered iPSC lines provided complementary platforms for evaluating the effects of constitutive and inducible ACE expression during granulocyte differentiation and function.

### Directed differentiation enables efficient generation of iPSC-derived neutrophils (iNeu)

To establish a scalable platform for granulocyte replacement therapy, we developed a directed differentiation protocol to efficiently generate human neutrophils from iPSCs. Differentiation was initiated through embryoid body (EB) formation under cytokine-free conditions to promote spontaneous germ layer specification. After 4 days in suspension culture, EBs were transferred to Matrigel-coated adherent conditions to support the emergence and expansion of hematopoietic progenitors. Subsequent culture in the presence of G-CSF drove terminal neutrophil differentiation (**Figure 1E**).

The differentiation protocol produced a highly enriched neutrophil population. By day 15, more than 80% of cells exhibited a mature neutrophil phenotype, as determined by CD11b and CD33 expression (**Figure 1F**). Notably, individual EB cultures continuously generated neutrophils over 5-6 collection cycles beginning approximately 14 days after differentiation initiation, yielding up to ∼1.5 × 10^7^ iPSC-derived neutrophils (iNeu) from 1 × 10^6^ input iPSCs. These findings demonstrate the platform’s robust expansion potential and scalability for producing clinically relevant numbers of neutrophils.

We next evaluated ACE expression in engineered iPSC-derived neutrophils. Neutrophils differentiated from the constitutive ACE-overexpressing iPSC line (ACE-iNeu) exhibited markedly increased ACE protein expression compared with control neutrophils (Cnt-iNeu), as demonstrated by immunoblotting (∼50-fold increase) and flow cytometric analyses (∼7-fold increase) (**Figures 1G and 1H**). Similarly, neutrophils generated from the inducible ACE-iPSC line (iACE-iNeu) displayed robust Dox-dependent ACE expression. Treatment with 1 μg/ml Dox resulted in pronounced ACE upregulation relative to both untreated iACE-iNeu cells and iCnt-iNeu cells, as confirmed by immunoblotting and flow cytometry (**Figures 1I and 1J**). Together, these results establish a scalable neutrophil differentiation platform and demonstrate efficient constitutive and inducible ACE expression in engineered iPSC-derived neutrophils.

### ACE-enhanced iPSC-derived neutrophils exhibit superior antimicrobial activity against drug-resistant bacterial pathogens

To determine whether ACE overexpression enhances antimicrobial activity, we first evaluated intracellular bacterial killing using methicillin-resistant *Staphylococcus aureus* (MRSA). Constitutive ACE overexpression significantly improved the bactericidal capacity of iPSC-derived neutrophils. Following infection at a multiplicity of infection (MOI) of 20, ACE-iNeu cells showed markedly greater bacterial clearance than control iNeu cells, resulting in 1.5-fold and 3.4-fold reductions in viable intracellular MRSA at 2 and 5 h post-infection, respectively (**Figure 2A**). We next investigated whether this enhanced antimicrobial activity translated into therapeutic efficacy in vivo. Neutropenia was induced in NSG mice by cyclophosphamide administration, followed by systemic MRSA challenge and adoptive transfer of iNeu cells. Twenty-four hours after transfusion, mice receiving ACE-iNeu cells exhibited significantly lower bacteremia than animals treated with control iNeu cells or PBS (**Figure 2B**). Consistent with the reduction in circulating bacteria, ACE-iNeu transfusion also markedly limited MRSA dissemination to peripheral organs, resulting in substantially lower bacterial burdens in the spleen, lung, and liver compared with both control groups (**Figures 2C-2E**). These findings demonstrate that ACE overexpression substantially enhances the ability of iNeu cells to control systemic bacterial infection and tissue dissemination in neutropenic hosts.

**Figure 2.**
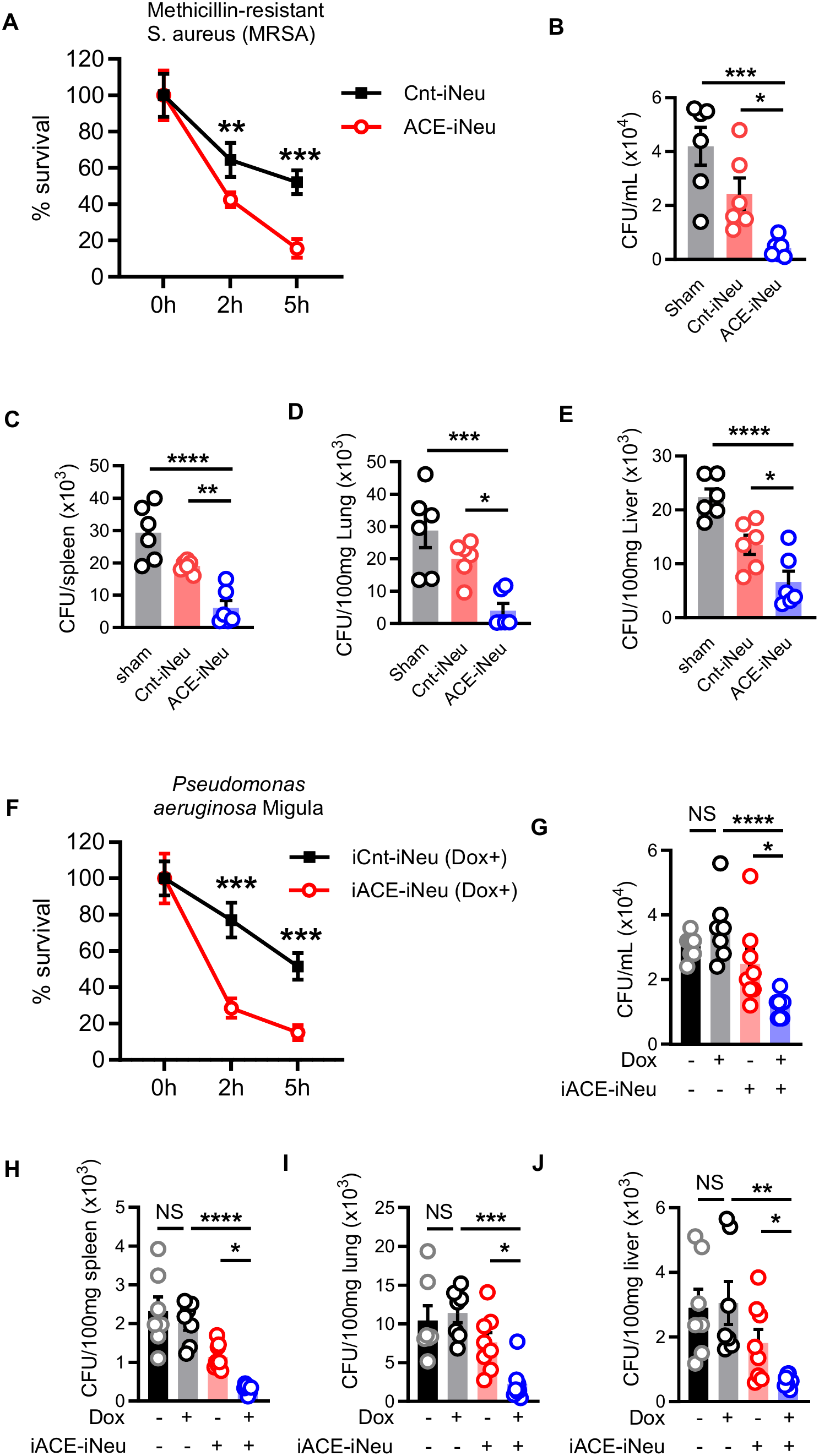
ACE-engineered iPSC-derived neutrophils enhance antimicrobial activity and improve bacterial clearance in neutropenic hosts. **(A)** In vitro intracellular killing of methicillin-resistant *Staphylococcus aureus* (MRSA) by ACE-iNeu and control iNeu (Cnt-iNeu) cells. Neutrophils were infected at a multiplicity of infection (MOI) of ∼20, and intracellular bacterial killing was quantified by colony-forming unit (CFU) assay. **(B–E)** In vivo bacterial clearance following transfusion of ACE-iNeu or Cont-iNeu cells into neutropenic NSG mice infected with MRSA. Bacterial burdens in blood (B), spleen (C), lungs (D), and liver (E) were quantified as indicated. **(F)** In vitro intracellular killing of multidrug-resistant *Pseudomonas aeruginosa* (Migula) by iACE-iNeu and iCnt-iNeu with Dox induction. Bacterial killing was assessed by CFU assay following infection at an MOI of ∼20. **(G–J)** In vivo bacterial clearance following transfusion of iACE-iNeu cells into neutropenic NSG mice infected with multidrug-resistant *P. aeruginosa*. Bacterial burdens in blood (G), spleen (H), lungs (I), and liver (J) were quantified as indicated. For *in vivo* studies, neutropenia was induced in NSG mice by a single intraperitoneal injection of cyclophosphamide (230 mg/kg). Twenty-four hours later, mice were intravenously challenged with either MRSA (**B–E**) or multidrug-resistant *P. aeruginosa* (**G–J**) (1 × 10^7^ CFU per mouse). One day after infection, mice received 1 × 10^7^ iNeu cells by tail-vein injection. MRSA-infected mice received ACE-iNeu, Cnt-iNeu, or PBS. *P. aeruginosa*-infected mice received iACE-iNeu or iCnt-iNeu. For inducible ACE expression, doxycycline (Dox) was administered in drinking water beginning 6 h before cell transfusion and maintained throughout the study. Blood bacterial counts were measured 24 h after transfusion, and bacterial burdens in spleen, lungs, and liver were determined 48 h after transfusion following PBS perfusion. Data are presented as mean ± SEM. Statistical significance was determined by one-way ANOVA with Bonferroni’s multiple-comparisons test. p < 0.05, p < 0.01, *p* < 0.001, p < 0.0001.

To test whether ACE augmentation could similarly enhance neutrophil activity against antibiotic-resistant pathogens, we next evaluated a multidrug-resistant *Pseudomonas aeruginosa* clinical isolate (BAA-2108) resistant to multiple antibiotics, including doxycycline. Consistent with its resistance profile, Dox had no detectable effect on bacterial growth, even at concentrations up to 5 μg/mL (**Figure S1**). We then assessed intracellular bacterial killing using Dox-inducible iACE-iNeu cells. Following ACE induction, iACE-iNeu cells exhibited significantly enhanced bactericidal activity, resulting in 1.5-fold and 1.9-fold reductions in viable intracellular bacteria at 2 and 5 h post-infection, respectively (**Figure 2F**). We next evaluated the therapeutic efficacy of iACE-iNeu transfusion in vivo using the neutropenic NSG infection model. Induction of ACE expression markedly improved bacterial clearance, resulting in significantly lower bacteremia compared with mice receiving non-induced iACE-iNeu cells or Dox alone (**Figure 2G**). Consistent with these findings, ACE induction significantly reduced bacterial burdens in the spleen, lung, and liver compared with both non-induced iACE-iNeu-treated mice and Dox-only controls (**Figures 2H-2J**). Collectively, these findings demonstrate that ACE induction enhances the antibacterial activity of iPSC-derived neutrophils and improves pathogen clearance both in vitro and in vivo, including against multidrug-resistant bacterial pathogens.

### ACE enhances oxidative burst and innate immune signaling in human neutrophils

Reactive oxygen species (ROS) production is a critical effector mechanism underlying neutrophil-mediated antimicrobial activity.^15^ To determine whether ACE regulates the oxidative response in human neutrophils, iACE-iNeu cells were stimulated with LPS (1 μg/ml) in the presence or absence of dox-induced ACE expression. Intracellular ROS generation was quantified using the dichlorodihydrofluorescein diacetate (DCFDA) assay and flow cytometric analysis. ACE induction significantly enhanced ROS production, resulting in a markedly greater oxidative burst compared with neutrophils expressing basal levels of ACE (**Figure 3A**).

**Figure 3.**
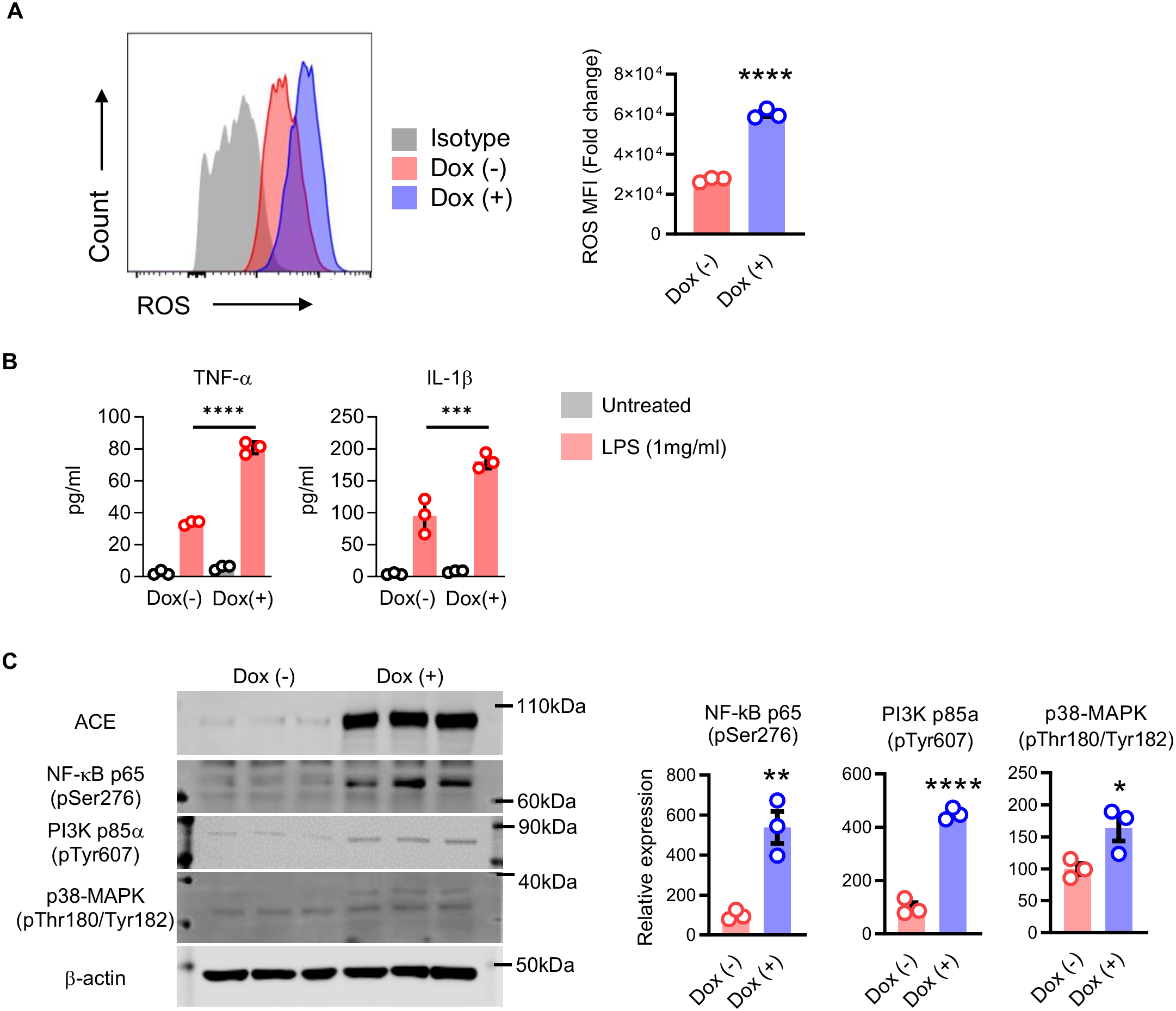
ACE potentiates inflammatory signaling and antimicrobial effector functions in human iPSC-derived neutrophils. **(A)** Intracellular reactive oxygen species (ROS) production in iACE-iNeu following stimulation with LPS (1 µg/ml, 30 min). ROS levels were measured by DCFDA staining and flow cytometry. Representative histograms are shown on the left, and quantification of mean fluorescence intensity (MFI) is shown on the right. **(B)** Cytokine production by iACE-iNeu following LPS stimulation. Cells were cultured in 6-well plates (3 × 10^6^ cells per well) and stimulated with LPS (500 ng/ml) for 12 h. Cytokine concentrations in culture supernatants were quantified by ELISA. **(C)** Immunoblot analysis of LPS-induced signaling pathway activation in iACE-iNeu. Phosphorylation of NF-κB p65, PI3K p85α, and p38 MAPK was evaluated following stimulation with LPS (500 ng/ml, 30 min). Band intensities were quantified using Odyssey software (v3.0), normalized to β-actin, and expressed relative to ACE-iNeu cells without ACE induction. For all experiments, ACE expression was induced by treatment with doxycycline (Dox; 1 µg/ml), whereas untreated iACE-iNeu served as controls. Data are presented as mean ± SEM from three independent experiments performed in triplicate. Statistical significance was determined using a two-tailed unpaired Student’s *t*-test for pairwise comparisons or one-way ANOVA with Bonferroni’s multiple-comparisons test, as appropriate. **p < 0.01, ***p < 0.001, ****p < 0.0001.

In addition to ROS generation, neutrophils contribute to host defense by secreting pro-inflammatory cytokines that amplify antimicrobial responses and coordinate the recruitment and activation of other immune cells.^16^ We therefore examined the production of the antibacterial cytokines IL-1β and TNFα following LPS stimulation. ACE induction significantly increased the production of both cytokines, with levels increasing approximately 2- to 3-fold relative to non-induced controls (**Figure 3B**). These findings indicate that ACE enhances not only the direct antimicrobial functions of neutrophils but also their capacity to orchestrate innate immune responses.

To further investigate the mechanisms underlying the enhanced antimicrobial activity of iACE-iNeu cells, we examined the PI3K, NF-κB, and p38 MAPK pathways, key regulators of neutrophil activation and innate immune responses.^12^ ACE induction significantly increased phosphorylation of PI3K, NF-κB, and p38 MAPK following LPS stimulation compared with non-induced controls (**Figure 3C**), consistent with enhanced activation of signaling pathways that regulate inflammatory and antimicrobial responses. Together, these findings demonstrate that ACE enhances oxidative burst, pro-inflammatory cytokine production, and innate immune signaling, thereby promoting the antimicrobial effector functions of human neutrophils.

### ACE promotes neutrophil maturation

Previous studies have implicated ACE in regulating hematopoiesis.^17–19^ To investigate its role in human myeloid development, we examined the effects of ACE induction during iPSC-directed myelopoiesis. iACE-iPSC cells were differentiated in the presence or absence of Dox, and hematopoietic progenitor and neutrophil populations were quantified throughout differentiation. Myeloid progenitors were isolated from EB cultures on day 10 and cultured in G-CSF-containing medium to support neutrophil development (**Figure S2**). ACE induction enhanced granulopoiesis and promoted progression through neutrophil developmental intermediates (**Figure 4A**). Dox-treated cultures generated substantially higher numbers of myeloid progenitors, pre-neutrophils, and mature neutrophils than non-induced cultures, whereas the immature neutrophil population was reduced. These findings suggest that ACE accelerates neutrophil maturation by facilitating progression through intermediate developmental stages and enhancing terminal differentiation. Collectively, these results demonstrate that ACE not only augments the antimicrobial effector functions of mature neutrophils but also promotes myeloid differentiation and increases neutrophil output.

**Figure 4.**
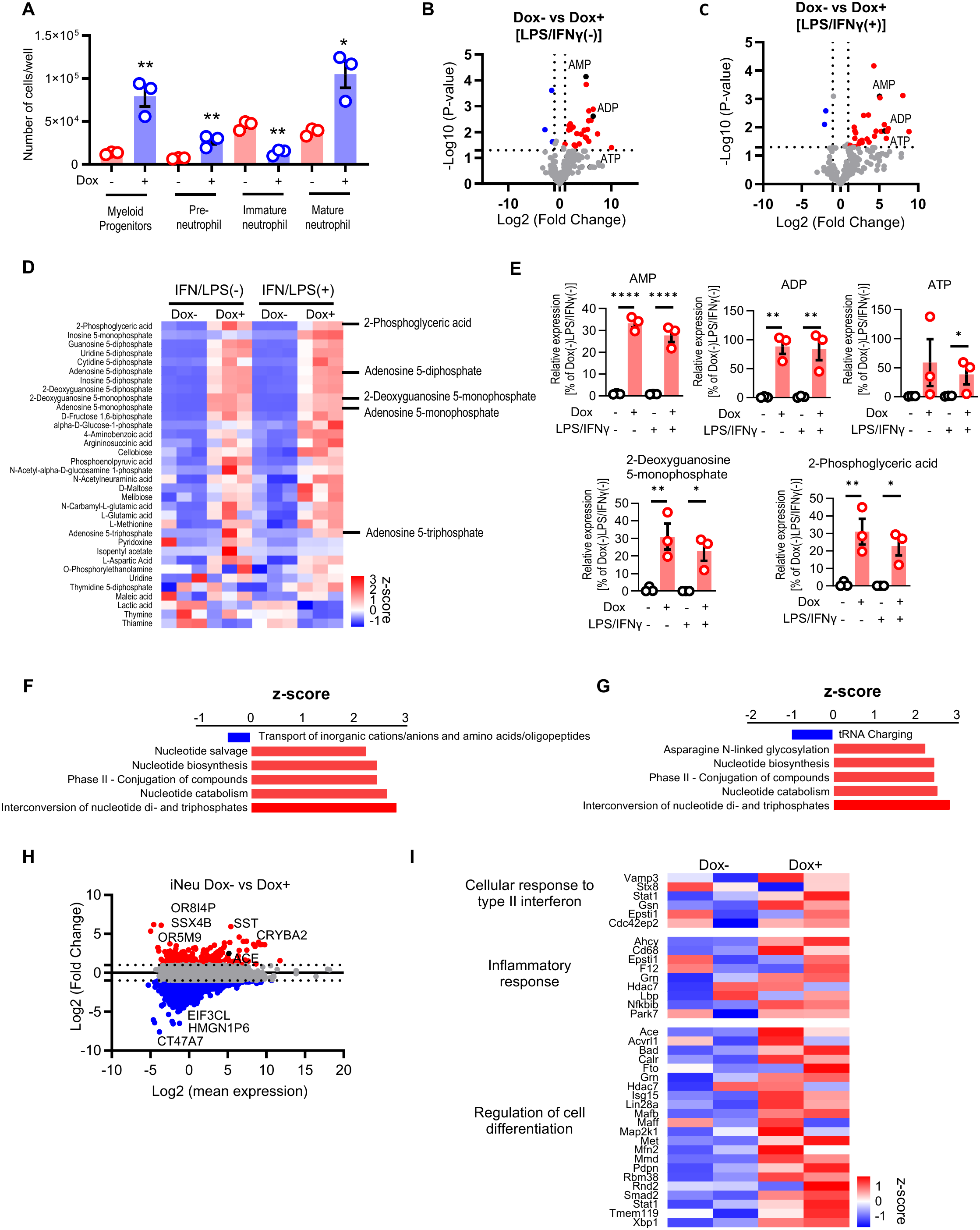
Transcriptomic and metabolomic profiling identifies ACE-dependent pathways associated with neutrophil maturation and function. iACE-iPSC lines were differentiated into myeloid progenitors and neutrophils in the presence or absence of doxycycline (Dox; 1 μg/ml) as described in the Methods. **(A)** Quantification of myeloid progenitors (CD66b^-^CD11b^-^CD34^+^), pre-neutrophils (CD66b^-^ CD11b^+^c-kit^+^), immature neutrophils (CD66b^-^CD11b^+^c-kit^-^), and mature neutrophils (CD66b^+^CD11b^+^) generated from iACE-iPSCs. Myeloid progenitors were harvested on day 10 of differentiation, seeded at 5 × 10^5^ cells per well in 24-well plates in RPMI supplemented with 100 ng/ml G-CSF, and analyzed by flow cytometry on day 15. Data are presented as mean ± SEM. Statistical significance was determined by two-way ANOVA with Bonferroni’s multiple-comparisons test. *p < 0.05, **p < 0.01. **(B and C)** Volcano plots of differentially abundant metabolites in Dox− versus Dox+ iACE-iNeu cells under basal conditions (B) or following LPS/IFNγ stimulation (C). Red dots indicate significantly increased metabolites. Differential metabolites were identified using a threshold of fold change >2 and p < 0.05. **(D)** Heatmap of significantly altered metabolites (p < 0.05) in Dox− versus Dox+ iACE-iNeu cells following LPS/IFNγ stimulation. Values are displayed as mean z-scores. **(E)** Relative intracellular levels of ADP, AMP, 2-deoxyguanosine 5′-monophosphate, and 2-phosphoglyceric acid in Dox− and Dox+ iACE-iNeu cells cultured with or without LPS/IFNγ stimulation. Metabolomic analyses were performed by mass spectrometry (n = 3 per group). **(F and G)** Ingenuity Pathway Analysis (IPA) of differentially abundant metabolites in Dox− versus Dox+ iACE-iNeu under basal conditions (F) or following LPS/IFNγ stimulation (G). Canonical pathways were ranked according to enrichment z-scores. **(H)** MA plot showing differentially expressed genes in Dox+ relative to Dox− iACE-iNeu cells. Red dots indicate significantly upregulated genes. Differential expression was defined as fold change >2. **(I)** Heatmap of differentially expressed genes associated with type II interferon responses, inflammatory responses, and regulation of cell differentiation in iACE-iNeu cells. RNA-seq, biologically independent samples per group.

### ACE reshapes metabolic and transcriptional programs during neutrophil differentiation

To investigate the mechanisms underlying ACE-mediated enhancement of neutrophil development and function, we performed untargeted mass spectrometry-based metabolomic profiling of iNeu cultured in the presence or absence of Dox-induced ACE expression. Under basal conditions, 29 metabolites were differentially abundant between ACE-induced and control iNeu (p < 0.01, fold change > 2.0), including 25 metabolites that were increased and 4 that were decreased following ACE induction (**Figure 4B**). Upon LPS/IFNγ stimulation, 30 metabolites were significantly altered, with 28 metabolites increased and 2 decreased in ACE-induced iNeu relative to controls (**Figure 4C**). Among the most prominently increased metabolites were ATP, ADP, and AMP, which were elevated 30- to 80-fold following ACE induction. For example, ATP was increased 59-fold under basal condition and 38-fold following LPS (**Figures 4D and 4E**). Consistent with these changes, Ingenuity Pathway Analysis (IPA) identified significant enrichment of pathways associated with nucleotide di- and triphosphate interconversion, suggesting enhanced bioenergetic capacity in ACE-expressing neutrophils (**Figures 4F and 4G**).

To determine how ACE influences transcriptional programs during human neutrophil development, we performed RNA sequencing of myeloid progenitors and mature iNeu generated from iACE-iPSCs differentiated in the presence or absence of Dox. Cell populations were purified by fluorescence-activated cell sorting prior to transcriptomic analysis. Gene set enrichment analysis revealed increased expression of Wnt signaling genes in both ACE-expressing myeloid progenitor and mature neutrophils (**Figure 4H and Figures S3A-S3C**).

Because metabolic remodeling is a key determinant of myeloid cell differentiation and function, we next examined pathways associated with cellular metabolism. ACE induction significantly enriched FXR/RXR and PPARα/RXRα signaling programs across both progenitor populations and mature neutrophils (**Figures S3D and S3E**). These findings suggest that ACE establishes metabolic programs during early granulopoiesis that persist throughout neutrophil maturation. Consistent with enhanced inflammatory responsiveness, Gene Ontology analysis of mature ACE-expressing neutrophils demonstrated enrichment of gene sets associated with the response to interferon-γ, inflammatory signaling, and cell differentiation (**Figure 4I**). Together, these data indicate that ACE coordinates metabolic and transcriptional programs that support neutrophil development and effector function.

## Discussion

Neutropenia remains a major cause of infection-related morbidity and mortality in patients undergoing chemotherapy and hematopoietic stem cell transplantation, particularly with the increasing prevalence of multidrug-resistant (MDR) pathogens.^20,21^ Although granulocyte transfusions provide temporary immune support, donor availability, variable cell yields, alloimmunization, and the short lifespan of mature neutrophils limit their clinical application.^22–25^ Here, we established a platform for generating iPSC-derived neutrophils and demonstrated that ACE engineering enhances neutrophil maturation, and antimicrobial activity. These findings identify ACE as a regulator of human neutrophil development and function and support the potential development of engineered granulocyte therapies.

Previous studies from our group demonstrated that ACE enhances innate immune responses in mice, whereas ACE deficiency compromises antibacterial defense.^11,26^ Extending these observations to human iPSC-derived neutrophils, we showed that constitutive and inducible ACE expression significantly improved intracellular killing of methicillin-resistant *Staphylococcus aureus* (MRSA) and MDR *Pseudomonas aeruginosa*. Importantly, adoptive transfer of ACE-engineered neutrophils into neutropenic mice reduced circulating bacterial burdens and dissemination to the spleen, lungs, and liver. These findings establish preclinical proof of concept for ACE-engineered neutrophils as a host-directed strategy against drug-resistant bacterial infections.

Mechanistically, ACE enhanced several antimicrobial effector pathways, including ROS generation, IL-1β and TNFα secretion, and activation of PI3K, NF-κB, and p38 MAPK signaling. These responses are consistent with our previous observations in murine immune cells and suggest that ACE coordinates multiple pathways involved in neutrophil activation and antibacterial defense rather than acting through a single downstream mechanism.^11,12^ Such coordinated enhancement may be particularly beneficial during severe infections requiring rapid and effective antimicrobial responses.

An important finding is the role of ACE in human granulopoiesis. ACE induction increased myeloid progenitor and mature neutrophil production while promoting progression through intermediate developmental stages. Although ACE has been implicated in hematopoietic regulation,^17–19^ its functional contribution to human neutrophil differentiation remains incompletely understood. Our findings suggest that ACE enhances both neutrophil generation and functional maturation, providing a dual advantage for developing engineered granulocyte products.

Transcriptomic profiling further revealed ACE-associated enrichment of developmental and inflammatory programs, including Wnt signaling, IFN-γ responses, and cell differentiation pathways. Wnt/β-catenin signaling is an established regulator of hematopoiesis and granulopoiesis,^27–33^ and similar ACE-associated transcriptional changes were previously observed in iPSC-derived macrophages.^34^ These findings suggest that ACE influences developmental programs contributing to neutrophil maturation, although the direct mechanistic relationship between ACE and Wnt signaling remains to be established.

Metabolomic analyses identified substantial ACE-dependent remodeling, including marked increases in ATP, ADP, and AMP under basal and inflammatory conditions, together with enrichment of nucleotide phosphate interconversion pathways. ACE expression was also associated with FXR/RXR and PPARα/RXRα signaling programs implicated in cellular metabolism and differentiation.^35^ These observations are consistent with previous studies demonstrating ACE-dependent regulation of cellular energy metabolism in macrophages and sperm.^13,36^ Given the importance of metabolic regulation in neutrophil development and activation,^37^ increased adenine nucleotide abundance may contribute to the enhanced oxidative burst, inflammatory signaling, and bactericidal activity observed in ACE-engineered neutrophils. However, further studies are needed to establish whether these metabolic changes directly mediate the functional effects of ACE.

Beyond its biological significance, our study has important translational implications. A major limitation of conventional granulocyte transfusion therapy is the inability to generate sufficient numbers of functionally optimized neutrophils in a standardized and donor-independent manner. The iPSC platform described here enables scalable neutrophil production while providing a genetically tractable system to enhance specific immune functions. The observation that ACE expression simultaneously promotes neutrophil production, survival-associated signaling, metabolic fitness, and antimicrobial activity highlights the potential value of engineering innate immune cells for therapeutic applications. Unlike conventional antimicrobial strategies that target microbial pathogens directly, ACE-enhanced neutrophils offer a host-directed approach that may remain effective even against highly drug-resistant organisms.

Collectively, our findings identify ACE as a central regulator of developmental, metabolic, and antimicrobial programs in human neutrophils. Through coordinated effects on granulopoiesis, cellular metabolism, inflammatory signaling, and effector function, ACE generates neutrophils with enhanced host-defense capacity. These results provide mechanistic insight into human neutrophil biology while laying the foundation for next-generation iPSC-derived granulocyte therapies. Future studies evaluating the persistence, safety, and efficacy of engineered neutrophils, including hypoimmunogenic universal donor products, may facilitate clinical translation of this strategy for the treatment of neutropenia and severe infections caused by multidrug-resistant pathogens.

## Acknowledgments

We thank Addgene (a non-profit organization) for providing the AAVS1_Puro_Tet3G_3xFLAG_Twin_Strep vector (#92099) used to construct the Tet-On-hACE plasmid, originally developed by Dr. Yannick Doyon’s Lab at the Centre Hospitalier Universitaire de Québec Research Center, Laval University, Quebec City, Canada. This work was supported by the Department of Pathology Mini-Grant and Startup Fund from Cedars-Sinai Medical Center, Los Angeles, CA, USA, awarded to Zakir Khan, and by National Institutes of Health (NIH) grant R01 AI164519 awarded to Kenneth E. Bernstein.

## Author Contributions

Conceptualization, Z.K., W.G.T., and K.E.B.; Methodology, T.S. and Z.K.; Investigation, T.S., S.A.B., S.S., T.B.D., B.G.C., E.A.B., N.K., and Z.K.; Formal Analysis, W.G.T., H.S.G., K.E.B., and Z.K.; Data Interpretation, D.S., W.G.T., H.S.G., K.E.B., and Z.K.; Visualization, T.S. and Z.K.; Writing – Original Draft, T.S. and Z.K.; Writing – Review & Editing, N.K., W.G.T., D.S., H.S.G., and K.E.B.; Supervision, Z.K. All authors reviewed and approved the final manuscript.

## Declaration of interests

A patent related to this study has been filed by Cedars-Sinai Medical Center on behalf of T.S., W.G.T., K.E.B., and Z.K. The authors declare that there is no conflict of interest regarding the publication of this article.

## Notes

### Competing Interest Statement

The authors have declared no competing interest.

