## Supplemental file for "ACE-engineered human iPSC-derived neutrophils enhance host defense against multidrug-resistant bacterial infections"

**This PDF file includes:**

Figures. S1 to S3
Tables S1
STAR Methods
Supplementary Text

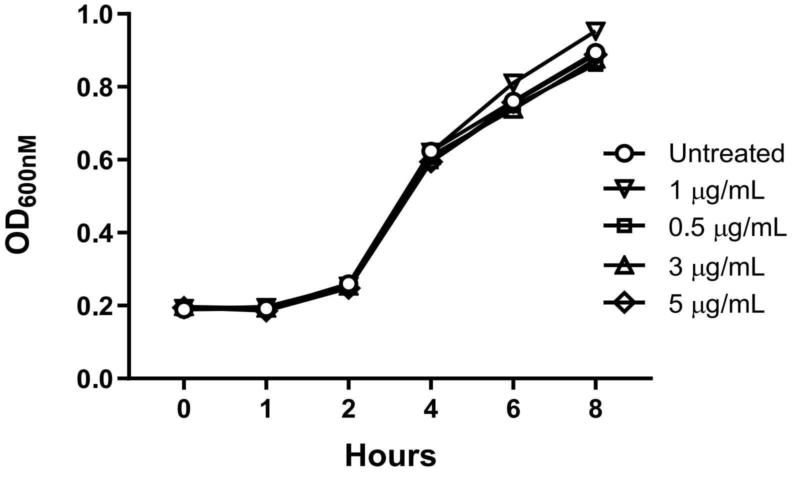


**Figure S1. Growth kinetics of *Pseudomonas aeruginosa* in the presence of doxycycline.** Overnight cultures of *Pseudomonas aeruginosa* (Migula strain) were diluted 1:300 into fresh LB medium containing doxycycline (Dox, 0-5 μg/ml). Bacterial growth was monitored by measuring optical density at 600 nm ($OD_{600}$) at the indicated time points to assess the direct effect of Dox on bacterial proliferation.

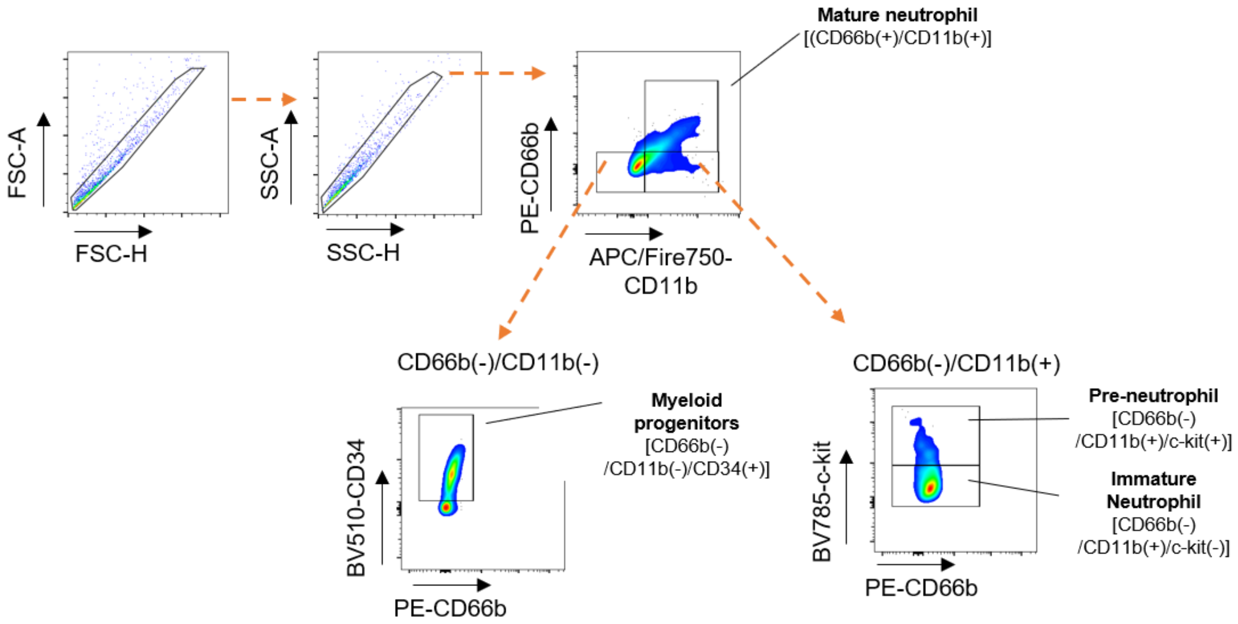


**Figure S2.** Flow cytometry gating strategy for identifying human iPSC-derived myeloid progenitors, pre-neutrophils, immature neutrophils, and mature neutrophils during differentiation.

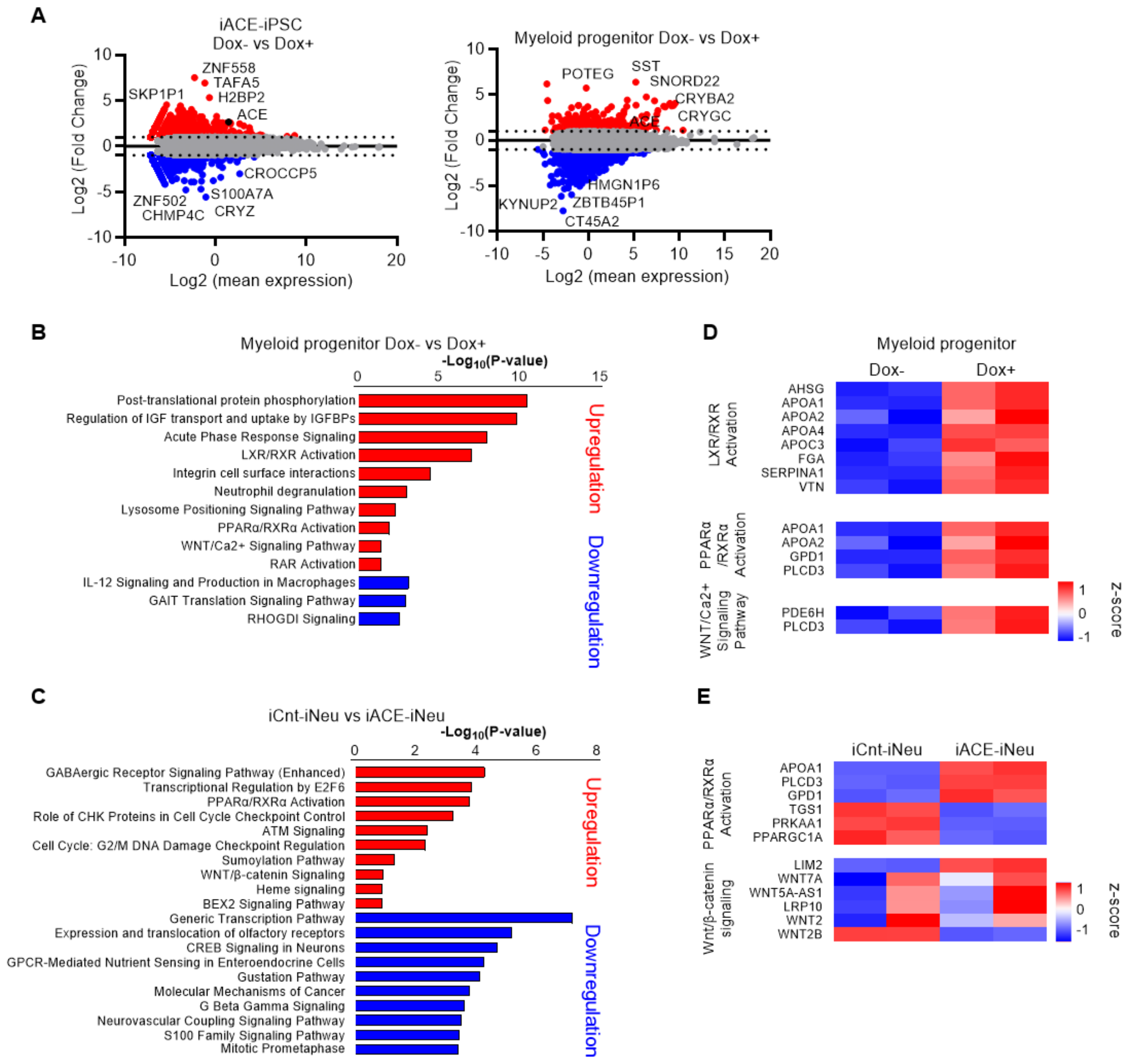


**Figure S3. ACE induces stage-specific transcriptional programs during myeloid differentiation of human iPSCs.**

ACE-iPSCs were differentiated into myeloid progenitors and neutrophils in the presence or absence of doxycycline (Dox; 1 μg/mL) as described in the Methods.

**(A)** MA plots showing differentially expressed genes in Dox-treated versus untreated iPSCs and myeloid progenitor. Red dots indicate significantly upregulated genes. Differential expression was defined as a fold change > 2.

**(B, C)** Ingenuity Pathway Analysis (IPA) of differentially expressed genes in iPSCs (B) and myeloid progenitor (C). Upregulated and downregulated gene sets were identified and ranked according to pathway enrichment.

**(D, E)** Heatmap analysis of genes associated with the PPARα/RXR and Wnt signaling pathway in myeloid progenitor (D) and iNeu cells (E).

**Table S1:** The list of flow cytometry antibodies.

| Antibodies | Source | Catalog Number |
|---|---|---|
| Human ACE | R&D | MAB9291-SP |
| APC/Fire™ 750 anti-mouse/human CD11b | Biolegend | 101262 |
| Brilliant Violet 785 anti-human CD117 (c-kit) | Biolegend | 313237 |
| Brilliant Violet 650 anti-human CD16 | Biolegend | 302041 |
| Brilliant Violet 510 anti-human CD34 | Biolegend | 343527 |
| Brilliant Violet 421™ anti-human CD33 | Biolegend | 303415 |
| PE anti-human CD66b | Biolegend | 305105 |

### Methods

#### Generation of constitutive and doxycycline-inducible human ACE expression constructs

To generate a constitutive human angiotensin-converting enzyme (hACE) expression construct, hACE cDNA was cloned into the NcoI restriction site of the AAVS1_Puro_Tet3G_3xFLAG_Twin_Strep vector (Addgene #92099), generating the hAAVS1 Tet3G-hACE-3xF-2xSTREP plasmid (unpublished data). The Tet-On promoter was subsequently replaced with the cytomegalovirus (CMV) promoter using the NsiI and AflII restriction sites, yielding the final constitutive expression construct, hAAVS1 CMV-hACE-3xF-2xSTREP. The complete plasmid sequence and annotation are provided in the Supplementary Materials.

For doxycycline (Dox)-inducible ACE expression, hACE cDNA was synthesized and inserted into the NcoI restriction site of the AAVS1_Puro_Tet3G_3xFLAG_Twin_Strep vector (Addgene #92099) under the control of the Tet-On 3G inducible expression system. The construct was generated by VectorBuilder (VectorBuilder ID: VB231104-1002hyg), and its complete sequence and annotation are provided in the Supplementary Materials.

For targeted genomic integration, each AAVS1-targeting construct was co-transfected with the Cas9-expression plasmid pXAT2 (Addgene #80494) to facilitate CRISPR-Cas9-mediated integration into the AAVS1 safe-harbor locus.

#### Human iPSC culture and directed differentiation into neutrophils

A previously established human induced pluripotent stem cell (hiPSC) line derived from human fibroblasts (CS88iCTR-nxx) was obtained from the iPSC Core at the Cedars-Sinai Biomanufacturing Center. Cells were maintained on Matrigel-coated culture plates (BD Biosciences, #35427) in mTeSR Plus medium (STEMCELL Technologies, #100-0276) at 37°C in a humidified atmosphere containing 5% $CO_2$. The culture medium was replaced every 2 days.

Mature neutrophils were generated from hiPSCs using an embryoid body (EB)-based differentiation protocol modified from previously published methods by Trump et al. and Lachmann et al.[1,2] Briefly, confluent hiPSCs cultured on Matrigel-coated plates (0.1 mg/mL) were maintained in EB medium consisting of Dulbecco's modified Eagle medium (DMEM) supplemented with 20% fetal bovine serum (FBS) and 1% nonessential amino acids (NEAA) for 3 days. Cells were then washed, treated with Versene solution (Gibco, #15040066) for 5 min, collected by centrifugation, and resuspended in EB medium. Subsequently, cells were seeded into 96-well ultra-low-attachment plates (Corning, #CLS3474) at a density of $2 \times 10^4$ cells/well to promote EB formation. After 4 days of suspension culture, 10 EBs were transferred into each well of Matrigel-coated 6-well plates and cultured in STEMdiff APEL2 medium (STEMCELL Technologies, #05275) supplemented with 25 ng/mL recombinant human interleukin-3 (hIL-3; PeproTech, #200-03) and 50 ng/mL recombinant human granulocyte colony-stimulating factor (hG-CSF; PeproTech, #300-25). Following attachment, EBs continuously released hematopoietic progenitor cells into the culture supernatant. Myeloid progenitors were collected from the supernatant every 3 days and subjected to terminal neutrophil differentiation by culturing for an additional 5 days in RPMI medium supplemented with 10% tetracycline-free FBS (Omega Scientific, #FB-15) and 100 ng/mL hG-CSF.

**Flow cytometry analysis**

Cells were stained with fluorophore-conjugated antibodies as listed in Table S1. For ACE detection, cells were incubated with a mouse anti-human ACE primary antibody (R&D Systems, #MAB9291; 1:100), followed by an Alexa Fluor 647-conjugated anti-mouse secondary antibody (Invitrogen, #A-21235; 1:500). Following immunostaining, cells were washed, resuspended in flow cytometry buffer, and analyzed using a Cytek NL-3000 flow cytometer. Flow cytometry data were analyzed using FlowJo software and are presented as representative histograms and mean fluorescence intensity (MFI), as appropriate.

**Western blot analysis**

Cells were washed twice with ice-cold phosphate-buffered saline (PBS) and lysed in radioimmunoprecipitation assay (RIPA) buffer supplemented with protease and phosphatase inhibitors (Thermo Fisher Scientific). Protein lysates were separated by sodium dodecyl sulfate–polyacrylamide gel electrophoresis (SDS-PAGE) and transferred onto polyvinylidene difluoride (PVDF) membranes. Membranes were incubated with primary antibodies against ACE (R&D Systems, #MAB9291; 1:1,000), β-actin (Sigma-Aldrich, #A3854; 1:1,000), phosphorylated NF-κB p65 (Novus Biologicals, #NB100-82086; 1:500), phosphorylated PI3K p85α (Cell Signaling Technology, #13666; 1:1,000), and phosphorylated p38 MAPK (Cell Signaling Technology, #9211; 1:1,000). Membranes were subsequently incubated with the appropriate infrared dye-conjugated secondary antibodies: IRDye 680RD goat anti-rabbit IgG (LI-COR Biosciences, #926-68071; 1:5,000), IRDye 800CW goat anti-mouse IgG (LI-COR Biosciences, #926-32210; 1:5,000), or IRDye 800CW donkey anti-goat IgG (LI-COR Biosciences, #926-32214; 1:5,000). Immunoreactive bands were visualized using an Odyssey CLx infrared imaging system (LI-COR Biosciences), and band intensities were quantified using Image Studio Lite software (version 5.2).

**Measurement of intracellular reactive oxygen species (ROS)**

To measure intracellular ROS production, $1 \times 10^6$ cells were stimulated with lipopolysaccharide (LPS; 1 μg/mL) for 30 min. Following stimulation, cells were resuspended in RPMI 1640 medium supplemented with 1% fetal bovine serum (FBS) and incubated with 20 μM 2′,7′-dichlorodihydrofluorescein diacetate (DCFDA; Abcam, #ab186028) for 30 min at 37°C. Cells were subsequently washed and stained with APC/Fire 750-conjugated anti-CD11b (BioLegend, #101262; 1:200) and Brilliant Violet 421-conjugated anti-CD33 (BioLegend, #303415; 1:100) antibodies for 30 min at 4°C. Following staining, cells were washed with fluorescence-activated cell sorting (FACS) buffer consisting of phosphate-buffered saline (PBS) supplemented with 2%

FBS, 0.1% sodium azide, and 1 mM EDTA. Intracellular ROS-associated fluorescence was measured using a Cytek NL-3000 flow cytometer.

**Measurement of cytokine production by ELISA**

Neutrophils ($3 \times 10^6$ cells/well) were stimulated with lipopolysaccharide (LPS; 500 ng/mL) for 12 h under the indicated experimental conditions, including Dox treatment (1 μg/mL) for inducible ACE-expressing cells, where applicable. Following incubation, culture supernatants were collected and stored at −80°C until analysis. The concentrations of interleukin-1β (IL-1β) and tumor necrosis factor-α (TNF-α) were quantified using commercially available enzyme-linked immunosorbent assay (ELISA) kits (IL-1β: Invitrogen, #BMS224-2; TNF-α: R&D Systems, #DY210-05), according to the manufacturers' instructions.

**Bacterial culture and quantification**

Two antibiotic-resistant bacterial strains were used in this study: methicillin-resistant *Staphylococcus aureus* (MRSA; LAC, USA300) and multidrug-resistant *Pseudomonas aeruginosa* (ATCC BAA-2108). MRSA was cultured in tryptic soy broth (TSB; BD Biosciences),[3] whereas *P. aeruginosa* was grown in Luria-Bertani (LB) broth (BD Biosciences).[4-7] Both bacterial strains were cultured overnight at 37°C with shaking. To obtain late-log-phase cultures, overnight cultures were diluted 1:100 in fresh medium and incubated for an additional 4 h at 37°C with shaking at 250 rpm. Bacteria were subsequently harvested by centrifugation at 4,000 rpm at 4°C, washed twice with phosphate-buffered saline (PBS), and resuspended in PBS. Bacterial concentrations were estimated by measuring optical density at 600 nm ($OD_{600}$), using an $OD_{600}$ of 0.3 as an approximate equivalent of $1 \times 10^8$ colony-forming units (CFU)/ml. The final bacterial inocula used in all in vitro and in vivo experiments were verified by serial dilution and colony enumeration on agar plates.

### In vitro intracellular bacterial killing assay

To evaluate intracellular bacterial killing activity, human iPSC-derived neutrophils (iNeu) were infected with either MRSA; LAC, USA300) or MRD *P. aeruginosa* at a multiplicity of infection (MOI) of approximately 20 for 30 min. Following infection, cells were treated with gentamicin (400 µg/ml) to eliminate extracellular bacteria. The completion of gentamicin treatment was designated as time zero (0 h). Intracellular bacterial survival was assessed at 0, 2, and 5 h by quantifying viable bacteria recovered from infected cells, expressed as colony-forming units (CFUs), as previously described.[4,8]

### Neutropenic mouse models of systemic bacterial infection and granulocyte transfusion

All animal experiments were approved by the Cedars-Sinai Institutional Animal Care and Use Committee (IACUC; protocol #010170). Immunodeficient NOD-scid IL2Rγnull (NSG) mice, aged 6–7 weeks, were obtained from The Jackson Laboratory (strain #002019). Mice were housed in microisolator cages under a 12-h light/dark cycle with ad libitum access to food and water. Animal health, including activity, feeding behavior, and signs of pain or distress, was monitored throughout the experiments in accordance with institutional animal care guidelines.

To evaluate the therapeutic efficacy of iPSC-derived neutrophil (iNeu) transfusion against systemic bacterial infections, mice were rendered neutropenic by a single intraperitoneal (i.p.) injection of cyclophosphamide (230 mg/kg), which induces neutropenia for approximately 4 days, as previously described.[4] Twenty-four hours after cyclophosphamide administration, mice were challenged systemically via retro-orbital injection with $1 \times 10^7$ CFUs of either MRSA or MRD *P. aeruginosa*. At 24 h post-infection, MRSA-infected mice received an intravenous (i.v.) transfusion of $1 \times 10^7$ constitutively ACE-expressing iNeu (ACE-iNeu), control iNeu (Cnt-iNeu), or phosphate-buffered saline (PBS; 50 µL) as a vehicle control. Mice infected with *P. aeruginosa* received an i.v. transfusion of $1 \times 10^7$ Dox-inducible ACE-expressing iNeu (iACE-iNeu). To induce ACE expression, Dox (150 μg/ml) was administered in the drinking water beginning 6 h before transfusion and maintained throughout the experiment.

Bacterial burdens in blood and tissues were evaluated sequentially using the same cohort of mice. At 24 h after granulocyte transfusion, peripheral blood samples were collected via retro-orbital bleeding to assess bacteremia. At 48 h after transfusion, the same mice were euthanized, and tissues were harvested to determine bacterial burdens. Harvested tissues were homogenized in 500 μL PBS, and bacterial counts in blood and tissue homogenates were quantified by serial dilution and plating on agar to determine viable CFUs, as previously described.[4,8]

**RNA sequencing and bioinformatics analysis**

Myeloid progenitors and mature neutrophils were isolated by fluorescence-activated cell sorting using a BD FACSAria III cell sorter (BD Biosciences). Sorted cells were washed twice with PBS, and total RNA was extracted using the RNeasy Plus Mini Kit (QIAGEN) according to the manufacturer's instructions. RNA samples with an A260/A280 ratio of 1.8–2.0 were selected for RNA sequencing (RNA-seq).

RNA sequencing was performed by BGI (https://www.bgi.com/global), and sequencing data were processed and analyzed using the Dr. Tom Multi-Omics Data Mining System (BGI; https://biosys.bgi.com/#/report/login).[9] Differentially expressed genes (DEGs) were identified using a significance threshold of $p < 0.05$ and a fold change > 2.0.

Functional enrichment and pathway analyses of DEGs were performed using Ingenuity Pathway Analysis (IPA; QIAGEN) and Gene Ontology (GO) analysis. For IPA, DEGs meeting the specified significance thresholds were mapped to canonical pathways and biological networks using the Ingenuity Pathway Knowledge Base, which integrates experimentally established and literature-derived gene functions and molecular interactions. GO enrichment analysis was performed using the Database for Annotation, Visualization and Integrated Discovery (DAVID;

https://david.ncifcrf.gov/), with functional annotations examined across three categories: biological process, molecular function, and cellular component.

For heatmap visualization, gene expression values expressed as transcripts per million (TPM) were $\log_2$-transformed, and gene-wise z-scores were calculated to display relative expression patterns across the analyzed samples.

**Metabolite profiling by liquid chromatography–tandem mass spectrometry (LC–MS/MS)**

Purified human iPSC-derived neutrophils (iNeu; $1 \times 10^7$ cells per sample) from doxycycline (Dox)-treated and untreated control groups were collected, washed, and subjected to metabolite extraction using 700 µl of ice-cold extraction solvent consisting of 40% acetonitrile, 40% methanol, and 20% water (v/v/v). Samples were vortexed vigorously for 5 min at 4°C and centrifuged at 10,000 × g for 10 min at 4°C. The resulting supernatants were collected and dried using a SpeedVac vacuum concentrator. Dried metabolite extracts were reconstituted by adding 20 µl methanol, followed by vortexing, and subsequently 80 µl water with additional vigorous vortexing. A 4-µl aliquot of each reconstituted sample was injected for liquid chromatography–tandem mass spectrometry (LC–MS/MS) analysis.

Metabolite profiling was performed as previously described.[10] Briefly, metabolite extracts were analyzed using an Agilent 6470A triple-quadrupole mass spectrometer coupled to an Agilent 1290 ultra-high-performance liquid chromatography (UHPLC) system (Agilent Technologies, Santa Clara, CA, USA). Mass spectrometric detection was performed in negative-ion mode. Chromatographic separation was achieved using an Agilent ZORBAX RRHD Extend-C18 column (2.1 × 150 mm, 1.8-µm particle size) equipped with a ZORBAX Extend-C18 Fast Guard column (2.1 mm, 1.8-µm particle size). Polar metabolites were analyzed using the Agilent MassHunter Metabolomics dynamic multiple reaction monitoring (dMRM) database and acquisition method. Chromatograms were processed and visualized using Agilent MassHunter Quantitative Analysis software for triple-quadrupole mass spectrometry. Integrated peak areas were manually reviewed to ensure consistent and accurate peak integration across samples.

**Supplementary Text**

**VB231104-1017 --- hAAVS1 CMV-hACE-3xF-2xSTREP Targeting Vector**
TGCTTTCTCTGACCAG – AAVS1 Arms
GAGGGCAGAGGA – T2A Sequence
ACCGAGTACAAGC – PuroR
AGACATGATAA – SV40 Poly A
GACATTGATTAT – CMV promoter + enhancer
ATGGGGGCC – hACE cDNA
CTGTGCCTT – bGH poly(A) signal
TTACCAATGC – AmpR

TCGCGCGTTTCGGTGATGACGGTGAAAACCTCTGACACATGCAGCTCCCGGAGACGGTCA
CAGCTTGTCTGTAAGCGGATGCCGGGAGCAGACAAGCCCGTCAGGGCGCGTCAGCGGGT
GTTGGCGGGTGTCGGGGCTGGCTTAACTATGCGGCATCAGAGCAGATTGTACTGAGAGTG
CACCATATGCGGTGTGAAATACCGCACAGATGCGTAAGGAGAAAATACCGCATCAGGCGC
CATTCGCCATTCAGGCTGCGCAACTGTTGGGAAGGGCGATCGGTGCGGGCCTCTTCGCTA
TTACGCCAGAATTCGCCCTTTGCTTTCTCTGACCAGCATTCTCTCCCCTGGGCCTGTGCCG
CTTTCTGTCTGCAGCTTGTGGCCTGGGTCACCTCTACGGCTGGCCCAGATCCTTCCCTGC
CGCCTCCTTCAGGTTCCGTCTTCCTCCACTCCCTCTTCCCCTTGCTCTCTGCTGTGTTGCT
GCCCAAGGATGCTCTTTCCGGAGCACTTCCTTCTCGGCGCTGCACCACGTGATGTCCTCT
GAGCGGATCCTCCCCGTGTCTGGGTCCTCTCCGGGCATCTCTCCTCCCTCACCCAACCCC
ATGCCGTCTTCACTCGCTGGGTTCCCTTTTCCTTCTCCTTCTGGGGCCTGTGCCATCTCTC
GTTTCTTAGGATGGCCTTCTCCGACGGATGTCTCCCTTGCGTCCCGCCTCCCCTTCTTGTA
GGCCTGCATCATCACCGTTTTTCTGGACAACCCCAAAGTACCCCGTCTCCCTGGCTTTAGC
CACCTCTCCATCCTCTTGCTTTCTTTGCCTGGACACCCCGTTCTCCTGTGGATTCGGGTCA
CCTCTCACTCCTTTCATTTGGGCAGCTCCCCTACCCCCCTTACCTCTCTAGTCTGTGCTAG
CTCTTCCAGCCCCCTGTCATGGCATCTTCCAGGGGTCCGAGAGCTCAGCTAGTCTTCTTCC
TCCAACCCGGGCCCCTATGTCCACTTCAGGACAGCATGTTTGCTGCCTCCAGGGATCCTG
TGTCCCCGAGCTGGGACCACCTTATATTCCCAGGGCCGGTTAATGTGGCTCTGGTTCTGG
GTACTTTTATCTGTCCCCTCCACCCCACAGTGGGGCAAGCTTCTGACCTCTTCTCTTCCTC
CCACAGGGCCTCGAGAGATCTGGCGGCGGAGAGGGCAGAGGAAGTCTTCTAACATGCGG
TGACGTGGAGGAGAATCCCGGCCCTAGCACCGAGTACAAGCCCACGGTGCGCCTCGCCA
CCCGCGACGACGTCCCCAGGGCCGTACGCACCCTCGCCGCCGCGTTCGCCGACTACCCC
GCCACGCGCCACACCGTCGATCCGGACCGCCACATCGAGCGGGTCACCGAGCTGCAAGA
ACTCTTCCTCACGCGCGTCGGGCTCGACATCGGCAAGGTGTGGGTCGCGGACGACGGCG
CCGCGGTGGCGGTCTGGACCACGCCGGAGAGCGTCGAAGCGGGGGCGGTGTTCGCCGA
GATCGGCCCGCGCATGGCCGAGTTGAGCGGTTCCCGGCTGGCCGCGCAGCAACAGATGG
AAGGCCTCCTGGCGCCGCACCGGCCCAAGGAGCCCGCGTGGTTCCTGGCCACCGTCGG
CGTCTCGCCCGACCACCAGGGCAAGGGTCTGGGCAGCGCCGTCGTGCTCCCCGGAGTG
GAGGCGGCCGAGCGCGCCGGGGTGCCCGCCTTCCTGGAGACCTCCGCGCCCCGCAACC
TCCCCTTCTACGAGCGGCTCGGCTTCACCGTCACCGCCGACGTCGAGGTGCCCGAAGGA
CCGCGCACCTGGTGCATGACCCGCAAGCCCGGTGCCTGATCTAGACAGACATGATAAGAT
ACATTGATGAGTTTGGACAAACCACAACTAGAATGCAGTGAAAAAAATGCTTTATTTGTGAA
ATTTGTGATGCTATTGCTTTATTTGTAACCATTATAAGCTGCAATAAACAAGTTAACAACAAC
AATTGCATTCATTTTATGTTTCAGGTTCAGGGGGAGGTGTGGGAGGTTTTTTAA**TGCAT**CCA
GATATACGCGTTGACATTGATTATTGACTAGTTATTAATAGTAATCAATTACGGGGTCATTA
GTTCATAGCCCATATATGGAGTTCCGCGTTACATAACTTACGGTAAATGGCCCGCCTGGCT
GACCGCCCAACGACCCCCGCCCATTGACGTCAATAATGACGTATGTTCCCATAGTAACGC
CAATAGGGACTTTCCATTGACGTCAATGGGTGGAGTATTTACGGTAAACTGCCCACTTGGC
AGTACATCAAGTGTATCATATGCCAAGTACGCCCCCTATTGACGTCAATGACGGTAAATGG

CCCGCCTGGCATTATGCCCAGTACATGACCTTATGGGACTTTCCTACTTGGCAGTACATCT
ACGTATTAGTCATCGCTATTACCATGGTGATGCGGTTTTGGCAGTACATCAATGGGCGTGG
ATAGCGGTTTGACTCACGGGGATTTCCAAGTCTCCACCCCATTGACGTCAATGGGAGTTTG
TTTTGGCACCAAAATCAACGGGACTTTCCAAAATGTCGTAACAACTCCGCCCCATTGACGC
AAATGGGCGGTAGGCGTGTACGGTGGGAGGTCTATATAAGCAGAGCTCTCTGGCTAACTA
GAGAACCCACTGCTTACTGGCTTATCGAAAC**CTTAAG**GCGGCCGCCGAGAATCTGTATTTT
CAGGGAGCGCCACCATGGGGGCCGCCTCGGGCCGCCGGGGGCCGGGGCTGCTGCTGC
CGCTGCCGCTGCTGTTGCTGCTGCCGCCGCAGCCCGCCCTGGCGTTGGACCACGGGCTG
CAGCCCGGCAACTTTTCTGCTGACGAGGCCGGGGCGCAGCTCTTCGCGCAGAGCTACAA
CTCCAGCGCCGAACAGGTGCTGTTCCAGAGCGTGGCCGCCAGCTGGGCGCACGACACCA
ACATCACCGCGGAGAATGCAAGGCGCCAGGAGGAAGCAGCCCTGCTCAGCCAGGAGTTT
GCGGAGGCCTGGGGCCAGAAGGCCAAGGAGCTGTATGAACCGATCTGGCAGAACTTCAC
GGACCCGCAGCTGCGCAGGATCATCGGAGCTGTGCGCACCCTGGGCTCTGCCAACCTGC
CCCTGGCTAAGCGGCAGCAGTACAACGCCCTGCTAAGCAACATGAGCAGGATCTACTCCA
CCGCCAAGGTCTGCCTCCCCAACAAGACTGCCACCTGCTGGTCCCTGGACCCAGATCTCA
CCAACATCCTGGCTTCCTCGCGAAGCTACGCCATGCTCCTGTTTGCCTGGGAGGGCTGGC
ACAACGCTGCGGGCATCCCGCTGAAACCGCTGTACGAGGATTTCACTGCCCTCAGCAATG
AAGCCTACAAGCAGGACGGCTTCACAGACACGGGGGCCTACTGGCGCTCCTGGTACAACT
CCCCCACCTTCGAGGACGATCTGGAACACCTCTACCAACAGCTAGAGCCCCTCTACCTGA
ACCTCCATGCCTTCGTCCGCCGCGCACTGCATCGCCGATACGGAGACAGATACATCAACC
TCAGGGGACCCATCCCTGCTCATCTGCTGGGAGACATGTGGGCCCAGAGCTGGGAAAACA
TCTACGACATGGTGGTGCCTTTCCCAGACAAGCCCAACCTCGATGTCACCAGTACTATGCT
GCAGCAGGGCTGGAACGCCACGCACATGTTCCGGGTGGCAGAGGAGTTCTTCACCTCCC
TGGAGCTCTCCCCCATGCCTCCCGAGTTCTGGGAAGGGTCGATGCTGGAGAAGCCGGCC
GACGGGCGGGAAGTGGTGTGCCACGCCTCGGCTTGGGACTTCTACAACAGGAAAGACTT
CAGGATCAAGCAGTGCACACGGGTCACGATGGACCAGCTCTCCACAGTGCACCATGAGAT
GGGCCATATACAGTACTACCTGCAGTACAAGGATCTGCCTGTCTCCCTGCGTCGGGGGGC
CAACCCCGGCTTCCATGAGGCCATTGGGGACGTGCTGGCGCTCTCGGTCTCCACTCCTGA
ACATCTGCACAAAATCGGCCTGCTGGACCGTGTCACCAATGACACGGAAAGTGACATCAAT
TACTTGCTAAAAATGGCACTGGAAAAAATTGCCTTCCTGCCCTTTGGCTACTTGGTGGACC
AGTGGCGCTGGGGGGTCTTTAGTGGGCGTACCCCCCCTTCCCGCTACAACTTCGACTGGT
GGTATCTTCGAACCAAGTATCAGGGGATCTGTCCTCCTGTTACCCGAAACGAAACCCACTT
TGATGCTGGAGCTAAGTTTCATGTTCCAAATGTGACACCATACATCAGGTACTTTGTGAGTT
TTGTCCTGCAGTTCCAGTTCCATGAAGCCCTGTGCAAGGAGGCAGGCTATGAGGGCCCAC
TGCACCAGTGTGACATCTACCGGTCCACCAAGGCAGGGGCCAAGCTCCGGAAGGTGCTG
CAGGCTGGCTCCTCCAGGCCCTGGCAGGAGGTGCTGAAGGACATGGTCGGCTTAGATGC
CCTGGATGCCCAGCCGCTGCTCAAGTACTTCCAGCTAGTCACCCTGTGGCTGCAGGAGCA
GAACCAGCAGAACGGCGAGGTCCTGGGCTGGCCCGAGTACCAGTGGCACCCGCCGTTGC
CTGACAACTACCCGGAGGGCATAGACCTGGTGACTGATGAGGCTGAGGCCAGCAAGTTTG
TGGAGGAATATGACCGGACATCCCAGGTGGTGTGGAACGAGTATGCCGAGGCCAACTGG
AACTACAACACCAACATCACCACAGAGACCAGCAAGATTCTGCTGCAGAAGAACATGCAAA
TAGCCAACCACACCCTAAAGTACGGCACCCAGGCCAGGAAGTTTGATGTGAACCAGTTGC
AGAACACCACTATCAAGCGGATCATAAAGAAGGTTCAGGACCTAGAACGGGCAGCACTGC
CTGCCCAGGAGCTGGAGGAGTACAACAAGATCCTGTTGGATATGGAAACCACCTACAGCG
TGGCCACTGTGTGCCACCCGAATGGCAGCTGCCTGCAGCTCGAGCCAGATCTGACGAATG
TGATGGCCACGTCCCGGAAATATGAAGACCTGTTATGGGCATGGGAGGGCTGGCGAGACA
AGGCGGGGAGAGCCATCCTCCAGTTTTACCCGAAATACGTGGAACTCATCAACCAGGCTG
CCCGGCTCAATGGCTATGTAGATGCAGGGGACTCGTGGAGGTCTATGTACGAGACACCAT
CCCTGGAGCAAGACCTGGAGCGGCTCTTCCAGGAGCTGCAGCCACTCTACCTCAACCTGC
ATGCCTACGTGCGCCGGGCCCTGCACCGTCACTACGGGGCCCAGCACATCAACCTGGAG
GGGCCCATTCCTGCTCACCTGCTGGGGAACATGTGGGCGCAGACCTGGTCCAACATCTAT

GACTTGGTGGTGCCCTTCCCTTCAGCCCCCTCGATGGACACCACAGAGGCTATGCTAAAG
CAGGGCTGGACGCCCAGGAGGATGTTTAAGGAGGCTGATGATTTCTTCACCTCCCTGGGG
CTGCTGCCCGTGCCTCCTGAGTTCTGGAACAAGTCGATGCTGGAGAAGCCAACCGACGGG
CGGGAGGTGGTCTGCCACGCCTCGGCCTGGGACTTCTACAACGGCAAGGACTTCCGGAT
CAAGCAGTGCACCACCGTGAACTTGGAGGACCTGGTGGTGGCCCACCACGAAATGGGCC
ACATCCAGTATTTCATGCAGTACAAAGACTTACCTGTGGCCTTGAGGGAGGGTGCCAACCC
CGGCTTCCATGAGGCCATTGGGGACGTGCTAGCCCTCTCAGTGTCTACGCCCAAGCACCT
GCACAGTCTCAACCTGCTGAGCAGTGAGGGTGGCAGCGACGAGCATGACATCAACTTTCT
GATGAAGATGGCCCTTGACAAGATCGCCTTTATCCCCTTCAGCTACCTCGTCGATCAGTGG
CGCTGGAGGGTATTTGATGGAAGCATCACCAAGGAGAACTATAACCAGGAGTGGTGGAGC
CTCAGGCTGAAGTACCAGGGCCTCTGCCCCCCAGTGCCCAGGACTCAAGGTGACTTTGAC
CCAGGGGCCAAGTTCCACATTCCTTCTAGCGTGCCTTACATCAGGTACTTTGTCAGCTTCA
TCATCCAGTTCCAGTTCCACGAGGCACTGTGCCAGGCAGCTGGCCACACGGGCCCCCTG
CACAAGTGTGACATCTACCAGTCCAAGGAGGCCGGGCAGCGCCTGGCGACCGCCATGAA
GCTGGGCTTCAGTAGGCCGTGGCCGGAAGCCATGCAGCTGATCACGGGCCAGCCCAACA
TGAGCGCCTCGGCCATGTTGAGCTACTTCAAGCCGCTGCTGGACTGGCTCCGCACGGAGA
ACGAGCTGCATGGGGAGAAGCTGGGCTGGCCGCAGTACAACTGGACGCCGAACTCCGCT
CGCTCAGAAGGGCCCCTCCCAGACAGCGGCCGCGTCAGCTTCCTGGGCCTGGACCTGGA
TGCGCAGCAGGCCCGCGTGGGCCAGTGGCTGCTGCTCTTCCTGGGCATCGCCCTGCTGG
TAGCCACCCTGGGCCTCAGCCAGCGGCTCTTCAGCATCCGCCACCGCAGCCTCCACCGG
CACTCCCACGGGCCCCAGTTCGGCTCCGAGGTGGAGCTGAGACACTCCGGATCCGCCGG
CGACTACAAGGACCACGACGGCGATTATAAGGATCACGACATCGACTACAAAGACGACGA
TGACAAGGGCGCCAGCAGCGCCTGGTCCCACCCTCAGTTTGAGAAGGGCGGAGGCTCTG
GCGGCGGAAGCGGAGGATCTGCTTGGAGCCACCCCCAGTTCGAAAAGTGATAACTCGAGT
CTAGACGTTTAAACCCTGCAGGCTGTGCCTTCTAGTTGCCAGCCATCTGTTGTTTGCCCCT
CCCCCGTGCCTTCCTTGACCCTGGAAGGTGCCACTCCCACTGTCCTTTCCTAATAAAATGA
GGAAATTGCATCGCATTGTCTGAGTAGGTGTCATTCTATTCTGGGGGGTGGGGTGGGGCA
GGACAGCAAGGGGGAGGATTGGGAAGACAATAGCAGGCATGCTGGGGATGTGGTGGGCT
CTATGGGGAGTACTAAGCTTTACTAGGGACAGGATTGGTGACAGAAAAGCCCCATCCTTAG
GCCTCCTCCTTCCTAGTCTCCTGATATTGGGTCTAACCCCCACCTCCTGTTAGGCAGATTC
CTTATCTGGTGACACACCCCCATTTCCTGGAGCCATCTCTCTCCTTGCCAGAACCTCTAAG
GTTTGCTTACGATGGAGCCAGAGAGGATCCTGGGAGGGAGAGCTTGGCAGGGGGTGGGA
GGGAAGGGGGGGATGCGTGACCTGCCCGGTTCTCAGTGGCCACCCTGCGCTACCCTCTC
CCAGAACCTGAGCTGCTCTGACGCGGCTGTCTGGTGCGTTTCACTGATCCTGGTGCTGCA
GCTTCCTTACACTTCCCAAGAGGAGAAGCAGTTTGGAAAAACAAAATCAGAATAAGTTGGT
CCTGAGTTCTAACTTTGGCTCTTCACCTTTCTAGTCCCCAATTTATATTGTTCCTCCGTGCG
TCAGTTTTACCTGTGAGATAAGGCCAGTAGCCAGCCCCGTCCTGGCAGGGCTGTGGTGAG
GAGGGGGGTGTCCGTGTGGAAAACTCCCTTTGTGAGAATGGTGCGTCCTAGGTGTTCACC
AGGTCGTGGCCGCCTCTACTCCCTTTCTCTTTCTCCATCCTTCTTTCCTTAAAGAGTCCCCA
GTGCTATCTGGGACATATTCCTCCGCCCAGAGCAGGGTCCCGCTTCCCTAAGGCCCTGCT
CTGGGCTTCTGGGTTTGAGTCCTTGGCAAGCCCAGGAGAGGCGCTCAGGCTTCCCTGTCC
CCCTTCCTCGTCCACCATCTCATGCCCCTGGCTCTCCTGCCCCTTCCCTACAGGGGTTCCT
GGCTCTGCTCTAAGGGCGAATTCTGCATTAATGAATCGGCCAACGCGCGGGGAGAGGCG
GTTTGCGTATTGGGCGCTCTTCCGCTTCCTCGCTCACTGACTCGCTGCGCTCGGTCGTTC
GGCTGCGGCGAGCGGTATCAGCTCACTCAAAGGCGGTAATACGGTTATCCACAGAATCAG
GGGATAACGCAGGAAAGAACATGTGAGCAAAAGGCCAGCAAAAGGCCAGGAACCGTAAAA
AGGCCGCGTTGCTGGCGTTTTTCCATAGGCTCCGCCCCCCTGACGAGCATCACAAAAATC
GACGCTCAAGTCAGAGGTGGCGAAACCCGACAGGACTATAAAGATACCAGGCGTTTCCCC
CTGGAAGCTCCCTCGTGCGCTCTCCTGTTCCGACCCTGCCGCTTACCGGATACCTGTCCG
CCTTTCTCCCTTCGGGAAGCGTGGCGCTTTCTCATAGCTCACGCTGTAGGTATCTCAGTTC
GGTGTAGGTCGTTCGCTCCAAGCTGGGCTGTGTGCACGAACCCCCCGTTCAGCCCGACC

GCTGCGCCTTATCCGGTAACTATCGTCTTGAGTCCAACCCGGTAAGACACGACTTATCGCC
ACTGGCAGCAGCCACTGGTAACAGGATTAGCAGAGCGAGGTATGTAGGCGGTGCTACAGA
GTTCTTGAAGTGGTGGCCTAACTACGGCTACACTAGAAGGACAGTATTTGGTATCTGCGCT
CTGCTGAAGCCAGTTACCTTCGGAAAAAGAGTTGGTAGCTCTTGATCCGGCAAACAAACCA
CCGCTGGTAGCGGTGGTTTTTTTGTTTGCAAGCAGCAGATTACGCGCAGAAAAAAAGGATC
TCAAGAAGATCCTTTGATCTTTTCTACGGGGTCTGACGCTCAGTGGAACGAAAACTCACGT
TAAGGGATTTTGGTCATGAGATTATCAAAAAGGATCTTCACCTAGATCCTTTTAAATTAAAAA
TGAAGTTTTAAATCAATCTAAAGTATATATGAGTAAACTTGGTCTGACAGTTACCAATGCTTA
ATCAGTGAGGCACCTATCTCAGCGATCTGTCTATTTCGTTCATCCATAGTTGCCTGACTCCC
CGTCGTGTAGATAACTACGATACGGGAGGGCTTACCATCTGGCCCCAGTGCTGCAATGAT
ACCGCGAGACCCACGCTCACCGGCTCCAGATTTATCAGCAATAAACCAGCCAGCCGGAAG
GGCCGAGCGCAGAAGTGGTCCTGCAACTTTATCCGCCTCCATCCAGTCTATTAATTGTTGC
CGGGAAGCTAGAGTAAGTAGTTCGCCAGTTAATAGTTTGCGCAACGTTGTTGCCATTGCTA
CAGGCATCGTGGTGTCACGCTCGTCGTTTGGTATGGCTTCATTCAGCTCCGGTTCCCAAC
GATCAAGGCGAGTTACATGATCCCCCATGTTGTGCAAAAAAGCGGTTAGCTCCTTCGGTCC
TCCGATCGTTGTCAGAAGTAAGTTGGCCGCAGTGTTATCACTCATGGTTATGGCAGCACTG
CATAATTCTCTTACTGTCATGCCATCCGTAAGATGCTTTTCTGTGACTGGTGAGTACTCAAC
CAAGTCATTCTGAGAATAGTGTATGCGGCGACCGAGTTGCTCTTGCCCGGCGTCAATACG
GGATAATACCGCGCCACATAGCAGAACTTTAAAAGTGCTCATCATTGGAAAACGTTCTTCG
GGGCGAAAACTCTCAAGGATCTTACCGCTGTTGAGATCCAGTTCGATGTAACCCACTCGTG
CACCCAACTGATCTTCAGCATCTTTTACTTTCACCAGCGTTTCTGGGTGAGCAAAAACAGG
AAGGCAAAATGCCGCAAAAAAGGGAATAAGGGCGACACGGAAATGTTGAATACTCATACTC
TTCCTTTTTCAATATTATTGAAGCATTTATCAGGGTTATTGTCTCATGAGCGGATACATATTT
GAATGTATTTAGAAAAATAAACAAATAGGGGTTCCGCGCACATTTCCCCGAAAAGTGCCAC
CTGACGTCTAAGAAACCATTATTATCATGACATTAACCTATAAAAATAGGCGTATCACGAGG
CCCTTTCGTC

**hAAVS1 Tet3G-hACE-3xF-2xSTREP Targeting Vector**

TGCTTTCTCTGACCAG – AAVS1 Arms
GAGGGCAGAGGA – T2A Sequence
ACCGAGTACAAGC – PuroR
AGACATGATAA – SV40 Poly A
TTACCCGGGGA – Tet-On 3G
GCGATCTGA – TRE3G BI Promoter
ATGGGGGCC – hACE cDNA
CTGTGCCTT – bGH poly(A) signal
TTACCAATGC – AmpR

TCGCGCGTTTCGGTGATGACGGTGAAAACCTCTGACACATGCAGCTCCCGGAGACGGTCA
CAGCTTGTCTGTAAGCGGATGCCGGGAGCAGACAAGCCCGTCAGGGCGCGTCAGCGGGT
GTTGGCGGGTGTCGGGGCTGGCTTAACTATGCGGCATCAGAGCAGATTGTACTGAGAGTG
CACCATATGCGGTGTGAAATACCGCACAGATGCGTAAGGAGAAAATACCGCATCAGGCGC
CATTCGCCATTCAGGCTGCGCAACTGTTGGGAAGGGCGATCGGTGCGGGCCTCTTCGCTA
TTACGCCAGAATTCGCCCTTTGCTTTCTCTGACCAGCATTCTCTCCCCTGGGCCTGTGCCG
CTTTCTGTCTGCAGCTTGTGGCCTGGGTCACCTCTACGGCTGGCCCAGATCCTTCCCTGC
CGCCTCCTTCAGGTTCCGTCTTCCTCCACTCCCTCTTCCCCTTGCTCTCTGCTGTGTTGCT
GCCCAAGGATGCTCTTTCCGGAGCACTTCCTTCTCGGCGCTGCACCACGTGATGTCCTCT
GAGCGGATCCTCCCCGTGTCTGGGTCCTCTCCGGGCATCTCTCCTCCCTCACCCAACCCC
ATGCCGTCTTCACTCGCTGGGTTCCCTTTTCCTTCTCCTTCTGGGGCCTGTGCCATCTCTC
GTTTCTTAGGATGGCCTTCTCCGACGGATGTCTCCCTTGCGTCCCGCCTCCCCTTCTTGTA
GGCCTGCATCATCACCGTTTTTCTGGACAACCCCAAAGTACCCCGTCTCCCTGGCTTTAGC
CACCTCTCCATCCTCTTGCTTTCTTTGCCTGGACACCCCGTTCTCCTGTGGATTCGGGTCA
CCTCTCACTCCTTTCATTTGGGCAGCTCCCCTACCCCCCTTACCTCTCTAGTCTGTGCTAG
CTCTTCCAGCCCCCTGTCATGGCATCTTCCAGGGGTCCGAGAGCTCAGCTAGTCTTCTTCC
TCCAACCCGGGCCCCTATGTCCACTTCAGGACAGCATGTTTGCTGCCTCCAGGGATCCTG
TGTCCCCGAGCTGGGACCACCTTATATTCCCAGGGCCGGTTAATGTGGCTCTGGTTCTGG
GTACTTTTATCTGTCCCCTCCACCCCACAGTGGGGCAAGCTTCTGACCTCTTCTCTTCCTC
CCACAGGGCCTCGAGAGATCTGGCGGCGGAGAGGGCAGAGGAAGTCTTCTAACATGCGG
TGACGTGGAGGAGAATCCCGGCCCTAGCACCGAGTACAAGCCCACGGTGCGCCTCGCCA
CCCGCGACGACGTCCCCAGGGCCGTACGCACCCTCGCCGCCGCGTTCGCCGACTACCCC
GCCACGCGCCACACCGTCGATCCGGACCGCCACATCGAGCGGGTCACCGAGCTGCAAGA
ACTCTTCCTCACGCGCGTCGGGCTCGACATCGGCAAGGTGTGGGTCGCGGACGACGGCG
CCGCGGTGGCGGTCTGGACCACGCCGGAGAGCGTCGAAGCGGGGGCGGTGTTCGCCGA
GATCGGCCCGCGCATGGCCGAGTTGAGCGGTTCCCGGCTGGCCGCGCAGCAACAGATGG
AAGGCCTCCTGGCGCCGCACCGGCCCAAGGAGCCCGCGTGGTTCCTGGCCACCGTCGG
CGTCTCGCCCGACCACCAGGGCAAGGGTCTGGGCAGCGCCGTCGTGCTCCCCGGAGTG
GAGGCGGCCGAGCGCGCCGGGGTGCCCGCCTTCCTGGAGACCTCCGCGCCCCGCAACC
TCCCCTTCTACGAGCGGCTCGGCTTCACCGTCACCGCCGACGTCGAGGTGCCCGAAGGA
CCGCGCACCTGGTGCATGACCCGCAAGCCCGGTGCCTGATCTAGACAGACATGATAAGAT
ACATTGATGAGTTTGGACAAACCACAACTAGAATGCAGTGAAAAAAATGCTTTATTTGTGAA
ATTTGTGATGCTATTGCTTTATTTGTAACCATTATAAGCTGCAATAAACAAGTTAACAACAAC
AATTGCATTCATTTTATGTTTCAGGTTCAGGGGGAGGTGTGGGAGGTTTTTTAATGCATTTA
CCCGGGGAGCATGTCAAGGTCAAAATCGTCAAGAGCGTCAGCAGGCAGCATATCAAGGTC
AAAGTCGTCAAGGGCATCGGCTGGGAGCATGTCTAAGTCAAAATCGTCAAGGGCGTCGGT
CGGCCCGCCGCTTTCGCACTTTAGCTGTTTCTCCAGGCCACATATGATTAGTTCCAGGCCG
AAAAGGAAGGCAGGTTCGGCTCCCTGCCGGTCGAACAGCTCAATTGCTTGTTTCAGAAGT
GGGGGCATAGAATCGGTGGTAGGTGTCTCTCTTTCCTCTTTTGCTACTTGATGCTCCTGTT
CCTCCAATACGCAGCCCAGTGTAAAGTGGCCCACGGCGGACAGAGCGTACAGTGCGTTCT

CCAGGGAGAAGCCTTGCTGACACAGGAACGCGAGCTGATTTTCCAGGGTTTCGTACTGTT
TCTCTGTTGGGCGGGTGCCGAGATGCACTTTAGCCCCGTCGCGATGTGAGAGGAGAGCA
CAGCGGTATGACTTGGCGTTGTTCCGCAGAAAGTCTTGCCATGACTCGCCTTCCAGGGGG
CAGGAGTGGGTATGATGCCTGTCCAGCATCTCGATTGGCAGGGCATCGAGCAGGGCCCG
CTTGTTCTTCACGTGCCAGTACAGGGTAGGCTGCTCAACTCCCAGCTTTTGAGCGAGTTTC
CTTGTCGTCAGGCCTTCGATACCGACTCCATTGAGTAATTCCAGAGCAGAGTTTATGACTTT
GCTCTTGTCCAGTCTAGACATGGTGAATTCTCCAGGCGATCTGACGGTTCACTAAACGAGC
TCTGCTTATATAGGCCTCCCACCGTACACGCCACCTCGACATACTCGAGTTTACTCCCTAT
CAGTGATAGAGAACGTATGAAGAGTTTACTCCCTATCAGTGATAGAGAACGTATGCAGACT
TTACTCCCTATCAGTGATAGAGAACGTATAAGGAGTTTACTCCCTATCAGTGATAGAGAACG
TATGACCAGTTTACTCCCTATCAGTGATAGAGAACGTATCTACAGTTTACTCCCTATCAGTG
ATAGAGAACGTATATCCAGTTTACTCCCTATCAGTGATAGAGAACGTATAAGCTTTAGGCGT
GTACGGTGGGCGCCTATAAAAGCAGAGCTCGTTTAGTGAACCGTCAGATCGCCTGGAGCA
ATTCCACAACACTTTTGTCTTATACCAACTTTCCGTACCACTTCCTACCCTCGTAAAGGTAC
CCTCAGCCTTAAGGCGGCCGCCGAGAATCTGTATTTTCAGGGAGCGCCACCATGGGGGC
CGCCTCGGGCCGCCGGGGGCCGGGGCTGCTGCTGCCGCTGCCGCTGCTGTTGCTGCTG
CCGCCGCAGCCCGCCCTGGCGTTGGACCACGGGCTGCAGCCCGGCAACTTTTCTGCTGA
CGAGGCCGGGGCGCAGCTCTTCGCGCAGAGCTACAACTCCAGCGCCGAACAGGTGCTGT
TCCAGAGCGTGGCCGCCAGCTGGGCGCACGACACCAACATCACCGCGGAGAATGCAAGG
CGCCAGGAGGAAGCAGCCCTGCTCAGCCAGGAGTTTGCGGAGGCCTGGGGCCAGAAGG
CCAAGGAGCTGTATGAACCGATCTGGCAGAACTTCACGGACCCGCAGCTGCGCAGGATCA
TCGGAGCTGTGCGCACCCTGGGCTCTGCCAACCTGCCCCTGGCTAAGCGGCAGCAGTAC
AACGCCCTGCTAAGCAACATGAGCAGGATCTACTCCACCGCCAAGGTCTGCCTCCCCAAC
AAGACTGCCACCTGCTGGTCCCTGGACCCAGATCTCACCAACATCCTGGCTTCCTCGCGA
AGCTACGCCATGCTCCTGTTTGCCTGGGAGGGCTGGCACAACGCTGCGGGCATCCCGCT
GAAACCGCTGTACGAGGATTTCACTGCCCTCAGCAATGAAGCCTACAAGCAGGACGGCTT
CACAGACACGGGGGCCTACTGGCGCTCCTGGTACAACTCCCCCACCTTCGAGGACGATCT
GGAACACCTCTACCAACAGCTAGAGCCCCTCTACCTGAACCTCCATGCCTTCGTCCGCCG
CGCACTGCATCGCCGATACGGAGACAGATACATCAACCTCAGGGGACCCATCCCTGCTCA
TCTGCTGGGAGACATGTGGGCCCAGAGCTGGGAAAACATCTACGACATGGTGGTGCCTTT
CCCAGACAAGCCCAACCTCGATGTCACCAGTACTATGCTGCAGCAGGGCTGGAACGCCAC
GCACATGTTCCGGGTGGCAGAGGAGTTCTTCACCTCCCTGGAGCTCTCCCCCATGCCTCC
CGAGTTCTGGGAAGGGTCGATGCTGGAGAAGCCGGCCGACGGGCGGGAAGTGGTGTGC
CACGCCTCGGCTTGGGACTTCTACAACAGGAAAGACTTCAGGATCAAGCAGTGCACACGG
GTCACGATGGACCAGCTCTCCACAGTGCACCATGAGATGGGCCATATACAGTACTACCTG
CAGTACAAGGATCTGCCTGTCTCCCTGCGTCGGGGGGCCAACCCCGGCTTCCATGAGGC
CATTGGGGACGTGCTGGCGCTCTCGGTCTCCACTCCTGAACATCTGCACAAAATCGGCCT
GCTGGACCGTGTCACCAATGACACGGAAAGTGACATCAATTACTTGCTAAAAATGGCACTG
GAAAAAATTGCCTTCCTGCCCTTTGGCTACTTGGTGGACCAGTGGCGCTGGGGGGTCTTT
AGTGGGCGTACCCCCCCTTCCCGCTACAACTTCGACTGGTGGTATCTTCGAACCAAGTATC
AGGGGATCTGTCCTCCTGTTACCCGAAACGAAACCCACTTTGATGCTGGAGCTAAGTTTCA
TGTTCCAAATGTGACACCATACATCAGGTACTTTGTGAGTTTTGTCCTGCAGTTCCAGTTCC
ATGAAGCCCTGTGCAAGGAGGCAGGCTATGAGGGCCCACTGCACCAGTGTGACATCTACC
GGTCCACCAAGGCAGGGGCCAAGCTCCGGAAGGTGCTGCAGGCTGGCTCCTCCAGGCCC
TGGCAGGAGGTGCTGAAGGACATGGTCGGCTTAGATGCCCTGGATGCCCAGCCGCTGCT
CAAGTACTTCCAGCTAGTCACCCTGTGGCTGCAGGAGCAGAACCAGCAGAACGGCGAGGT
CCTGGGCTGGCCCGAGTACCAGTGGCACCCGCCGTTGCCTGACAACTACCCGGAGGGCA
TAGACCTGGTGACTGATGAGGCTGAGGCCAGCAAGTTTGTGGAGGAATATGACCGGACAT
CCCAGGTGGTGTGGAACGAGTATGCCGAGGCCAACTGGAACTACAACACCAACATCACCA
CAGAGACCAGCAAGATTCTGCTGCAGAAGAACATGCAAATAGCCAACCACACCCTAAAGTA
CGGCACCCAGGCCAGGAAGTTTGATGTGAACCAGTTGCAGAACACCACTATCAAGCGGAT

CATAAAGAAGGTTCAGGACCTAGAACGGGCAGCACTGCCTGCCCAGGAGCTGGAGGAGT
ACAACAAGATCCTGTTGGATATGGAAACCACCTACAGCGTGGCCACTGTGTGCCACCCGA
ATGGCAGCTGCCTGCAGCTCGAGCCAGATCTGACGAATGTGATGGCCACGTCCCGGAAAT
ATGAAGACCTGTTATGGGCATGGGAGGGCTGGCGAGACAAGGCGGGGAGAGCCATCCTC
CAGTTTTACCCGAAATACGTGGAACTCATCAACCAGGCTGCCCGGCTCAATGGCTATGTAG
ATGCAGGGGACTCGTGGAGGTCTATGTACGAGACACCATCCCTGGAGCAAGACCTGGAGC
GGCTCTTCCAGGAGCTGCAGCCACTCTACCTCAACCTGCATGCCTACGTGCGCCGGGCCC
TGCACCGTCACTACGGGGCCCAGCACATCAACCTGGAGGGGCCCATTCCTGCTCACCTGC
TGGGGAACATGTGGGCGCAGACCTGGTCCAACATCTATGACTTGGTGGTGCCCTTCCCTT
CAGCCCCCTCGATGGACACCACAGAGGCTATGCTAAAGCAGGGCTGGACGCCCAGGAGG
ATGTTTAAGGAGGCTGATGATTTCTTCACCTCCCTGGGGCTGCTGCCCGTGCCTCCTGAGT
TCTGGAACAAGTCGATGCTGGAGAAGCCAACCGACGGGCGGGAGGTGGTCTGCCACGCC
TCGGCCTGGGACTTCTACAACGGCAAGGACTTCCGGATCAAGCAGTGCACCACCGTGAAC
TTGGAGGACCTGGTGGTGGCCCACCACGAAATGGGCCACATCCAGTATTTCATGCAGTAC
AAAGACTTACCTGTGGCCTTGAGGGAGGGTGCCAACCCCGGCTTCCATGAGGCCATTGGG
GACGTGCTAGCCCTCTCAGTGTCTACGCCCAAGCACCTGCACAGTCTCAACCTGCTGAGC
AGTGAGGGTGGCAGCGACGAGCATGACATCAACTTTCTGATGAAGATGGCCCTTGACAAG
ATCGCCTTTATCCCCTTCAGCTACCTCGTCGATCAGTGGCGCTGGAGGGTATTTGATGGAA
GCATCACCAAGGAGAACTATAACCAGGAGTGGTGGAGCCTCAGGCTGAAGTACCAGGGCC
TCTGCCCCCCAGTGCCCAGGACTCAAGGTGACTTTGACCCAGGGGCCAAGTTCCACATTC
CTTCTAGCGTGCCTTACATCAGGTACTTTGTCAGCTTCATCATCCAGTTCCAGTTCCACGAG
GCACTGTGCCAGGCAGCTGGCCACACGGGCCCCCTGCACAAGTGTGACATCTACCAGTC
CAAGGAGGCCGGGCAGCGCCTGGCGACCGCCATGAAGCTGGGCTTCAGTAGGCCGTGG
CCGGAAGCCATGCAGCTGATCACGGGCCAGCCCAACATGAGCGCCTCGGCCATGTTGAG
CTACTTCAAGCCGCTGCTGGACTGGCTCCGCACGGAGAACGAGCTGCATGGGGAGAAGC
TGGGCTGGCCGCAGTACAACTGGACGCCGAACTCCGCTCGCTCAGAAGGGCCCCTCCCA
GACAGCGGCCGCGTCAGCTTCCTGGGCCTGGACCTGGATGCGCAGCAGGCCCGCGTGG
GCCAGTGGCTGCTGCTCTTCCTGGGCATCGCCCTGCTGGTAGCCACCCTGGGCCTCAGC
CAGCGGCTCTTCAGCATCCGCCACCGCAGCCTCCACCGGCACTCCCACGGGCCCCAGTT
CGGCTCCGAGGTGGAGCTGAGACACTCCGGATCCGCCGGCGACTACAAGGACCACGACG
GCGATTATAAGGATCACGACATCGACTACAAAGACGACGATGACAAGGGCGCCAGCAGCG
CCTGGTCCCACCCTCAGTTTGAGAAGGGCGGAGGCTCTGGCGGCGGAAGCGGAGGATCT
GCTTGGAGCCACCCCCAGTTCGAAAAGTGATAACTCGAGTCTAGACGTTTAAACCCTGCAG
GCTGTGCCTTCTAGTTGCCAGCCATCTGTTGTTTGCCCCTCCCCCGTGCCTTCCTTGACCC
TGGAAGGTGCCACTCCCACTGTCCTTTCCTAATAAAATGAGGAAATTGCATCGCATTGTCT
GAGTAGGTGTCATTCTATTCTGGGGGGTGGGGTGGGGCAGGACAGCAAGGGGGAGGATT
GGGAAGACAATAGCAGGCATGCTGGGGATGTGGTGGGCTCTATGGGGAGTACTAAGCTTT
ACTAGGGACAGGATTGGTGACAGAAAAGCCCCATCCTTAGGCCTCCTCCTTCCTAGTCTCC
TGATATTGGGTCTAACCCCCACCTCCTGTTAGGCAGATTCCTTATCTGGTGACACACCCCC
ATTTCCTGGAGCCATCTCTCTCCTTGCCAGAACCTCTAAGGTTTGCTTACGATGGAGCCAG
AGAGGATCCTGGGAGGGAGAGCTTGGCAGGGGGTGGGAGGGAAGGGGGGGATGCGTGA
CCTGCCCGGTTCTCAGTGGCCACCCTGCGCTACCCTCTCCCAGAACCTGAGCTGCTCTGA
CGCGGCTGTCTGGTGCGTTTCACTGATCCTGGTGCTGCAGCTTCCTTACACTTCCCAAGAG
GAGAAGCAGTTTGGAAAAACAAAATCAGAATAAGTTGGTCCTGAGTTCTAACTTTGGCTCTT
CACCTTTCTAGTCCCCAATTTATATTGTTCCTCCGTGCGTCAGTTTTACCTGTGAGATAAGG
CCAGTAGCCAGCCCCGTCCTGGCAGGGCTGTGGTGAGGAGGGGGGTGTCCGTGTGGAAA
ACTCCCTTTGTGAGAATGGTGCGTCCTAGGTGTTCACCAGGTCGTGGCCGCCTCTACTCC
CTTTCTCTTTCTCCATCCTTCTTTCCTTAAAGAGTCCCCAGTGCTATCTGGGACATATTCCT
CCGCCCAGAGCAGGGTCCCGCTTCCCTAAGGCCCTGCTCTGGGCTTCTGGGTTTGAGTCC
TTGGCAAGCCCAGGAGAGGCGCTCAGGCTTCCCTGTCCCCCTTCCTCGTCCACCATCTCA
TGCCCCTGGCTCTCCTGCCCCTTCCCTACAGGGGTTCCTGGCTCTGCTCTAAGGGCGAAT

TCTGCATTAATGAATCGGCCAACGCGCGGGGAGAGGCGGTTTGCGTATTGGGCGCTCTTC
CGCTTCCTCGCTCACTGACTCGCTGCGCTCGGTCGTTCGGCTGCGGCGAGCGGTATCAG
CTCACTCAAAGGCGGTAATACGGTTATCCACAGAATCAGGGGATAACGCAGGAAAGAACAT
GTGAGCAAAAGGCCAGCAAAAGGCCAGGAACCGTAAAAAGGCCGCGTTGCTGGCGTTTTT
CCATAGGCTCCGCCCCCCTGACGAGCATCACAAAAATCGACGCTCAAGTCAGAGGTGGCG
AAACCCGACAGGACTATAAAGATACCAGGCGTTTCCCCCTGGAAGCTCCCTCGTGCGCTC
TCCTGTTCCGACCCTGCCGCTTACCGGATACCTGTCCGCCTTTCTCCCTTCGGGAAGCGT
GGCGCTTTCTCATAGCTCACGCTGTAGGTATCTCAGTTCGGTGTAGGTCGTTCGCTCCAAG
CTGGGCTGTGTGCACGAACCCCCCGTTCAGCCCGACCGCTGCGCCTTATCCGGTAACTAT
CGTCTTGAGTCCAACCCGGTAAGACACGACTTATCGCCACTGGCAGCAGCCACTGGTAAC
AGGATTAGCAGAGCGAGGTATGTAGGCGGTGCTACAGAGTTCTTGAAGTGGTGGCCTAAC
TACGGCTACACTAGAAGGACAGTATTTGGTATCTGCGCTCTGCTGAAGCCAGTTACCTTCG
GAAAAAGAGTTGGTAGCTCTTGATCCGGCAAACAAACCACCGCTGGTAGCGGTGGTTTTT
TGTTTGCAAGCAGCAGATTACGCGCAGAAAAAAAGGATCTCAAGAAGATCCTTTGATCTTTT
CTACGGGGTCTGACGCTCAGTGGAACGAAAACTCACGTTAAGGGATTTTGGTCATGAGATT
ATCAAAAAGGATCTTCACCTAGATCCTTTTAAATTAAAAATGAAGTTTTAAATCAATCTAAAG
TATATATGAGTAAACTTGGTCTGACAGTTACCAATGCTTAATCAGTGAGGCACCTATCTCAG
CGATCTGTCTATTTCGTTCATCCATAGTTGCCTGACTCCCCGTCGTGTAGATAACTACGATA
CGGGAGGGCTTACCATCTGGCCCCAGTGCTGCAATGATACCGCGAGACCCACGCTCACC
GGCTCCAGATTTATCAGCAATAAACCAGCCAGCCGGAAGGGCCGAGCGCAGAAGTGGTCC
TGCAACTTTATCCGCCTCCATCCAGTCTATTAATTGTTGCCGGGAAGCTAGAGTAAGTAGTT
CGCCAGTTAATAGTTTGCGCAACGTTGTTGCCATTGCTACAGGCATCGTGGTGTCACGCTC
GTCGTTTGGTATGGCTTCATTCAGCTCCGGTTCCCAACGATCAAGGCGAGTTACATGATCC
CCCATGTTGTGCAAAAAAGCGGTTAGCTCCTTCGGTCCTCCGATCGTTGTCAGAAGTAAGT
TGGCCGCAGTGTTATCACTCATGGTTATGGCAGCACTGCATAATTCTCTTACTGTCATGCC
ATCCGTAAGATGCTTTTCTGTGACTGGTGAGTACTCAACCAAGTCATTCTGAGAATAGTGTA
TGCGGCGACCGAGTTGCTCTTGCCCGGCGTCAATACGGGATAATACCGCGCCACATAGCA
GAACTTTAAAAGTGCTCATCATTGGAAAACGTTCTTCGGGGCGAAAACTCTCAAGGATCTTA
CCGCTGTTGAGATCCAGTTCGATGTAACCCACTCGTGCACCCAACTGATCTTCAGCATCTT
TTACTTTCACCAGCGTTTCTGGGTGAGCAAAAACAGGAAGGCAAAATGCCGCAAAAAAGGG
AATAAGGGCGACACGGAAATGTTGAATACTCATACTCTTCCTTTTTCAATATTATTGAAGCA
TTTATCAGGGTTATTGTCTCATGAGCGGATACATATTTGAATGTATTTAGAAAAATAAACAAA
TAGGGGTTCCGCGCACATTTCCCCGAAAAGTGCCACCTGACGTCTAAGAAACCATTATTAT
CATGACATTAACCTATAAAAATAGGCGTATCACGAGGCCCTTTCGTC
